# Characterizing the role of exosomal miRNAs in metastasis

**DOI:** 10.1101/2024.08.20.608894

**Authors:** Piyush Agrawal, Gulden Olgun, Arashdeep Singh, Vishaka Gopalan, idhar Hannenhalli

**Author notes:** **Co-corresponding Author’s:** 1. **Piyush Agrawal,** Department of Medical Research, SRM Medical College Hospital & Research Centre, SRMIST, Kattankulathur, Chennai, Tamil Nadu, India, 2. **Sridhar Hannenhalli,** National Cancer Institute, Bethesda, MD, USA.

## Abstract

**Background:** Exosomal microRNAs (exomiRs), transported via exosomes, play a pivotal role in intercellular communication. In cancer, exomiRs influence tumor progression by regulating key cellular processes such as proliferation, angiogenesis, and metastasis. Their role in mediating communication between cancer cells and the tumor microenvironment highlights their significance as potential diagnostic and therapeutic targets.

**Methodology:** In this study, we aimed to characterize the role of exomiRs in influencing the pre-metastatic niche (PMN). Across 7 tumor types, including 4 cell lines and three tumors, we extracted high confidence exomiRs (Log FC >= 2 in exosomes relative to control) and their targets (experimentally identified and targeted by at least 2 exomiRs). Subsequently, we identified enriched pathways and selected the top 100 high-confidence exomiR targets based on the frequency of their appearance in the enriched pathways. These top 100 targets were consistently used throughout the analysis.

**Results:** Cancer cell line and tumor derived ExomiRs have significantly higher GC content relative to genomic background. Pathway enriched among the top exomiR targets included general cancer-associated processes such as “wound healing” and “regulation of epithelial cell proliferation”, as well as cancer-specific processes, such as “regulation of angiogenesis in kidney” (KIRC), “ossification” in lung (LUAD), and “positive regulation of cytokine production” in pancreatic cancer (PAAD). Similarly, ‘Pathways in cancer’ and ‘MicroRNAs in cancer’ ranked among the top 10 enriched KEGG pathways in all cancer types. ExomiR targets were not only enriched for cancer-specific tumor suppressor genes (TSG) but are also downregulated in pre-metastatic niche formed in lungs compared to normal lung. Motif analysis shows high similarity among motifs identified from exomiRs across cancer types. Our analysis recapitulates exomiRs associated with M2 macrophage differentiation and chemoresistance such as miR-21 and miR-222-3p, regulating signaling pathways such as PTEN/PI3/Akt, NF-κB, etc. Cox regression indicated that exomiR targets are significantly associated with overall survival of patients in TCGA. Lastly, a Support Vector Machine (SVM) model using exomiR target gene expression classified responders and non-responders to neoadjuvant chemotherapy with an AUROC of 0.96 (in LUAD), higher than other previously reported gene signatures.

**Conclusion:** Our study characterizes the pivotal role of exomiRs in shaping the PMN in diverse cancers, underscoring their diagnostic and therapeutic potential.

## 1. Introduction

Exosomes belong to a group of small membrane vesicles formed by the inward budding of endosomal membranes, which eventually fuse with the plasma membrane to secrete exosomes into the extracellular environment [1]. Exosomes play a crucial role in intercellular communication by facilitating transfer of various bioactive molecules such as DNA, RNA, protein, lipids, etc. which can affect the recipient cell’s function [2]. In normal conditions, exosomes are involved in immune response, tissue repair, and cellular homeostasis. However, in diseased conditions such as cancer, they can contribute to cancer progression by transferring specific biomolecules, in particular, miRNAs, to promote angiogenesis and metastasis [3,4].

Metastasis is the primary cause of cancer-related mortality [5]. Exosomes, via its cargo, play an important role in cancer metastasis by mediating intercellular communication and preparing distant sites for tumor cell colonization, also known as pre-metastatic niche (PMN) formation [6,7]. They also modulate the immune response by carrying various immunosuppressive molecules, enabling cancer cells evade immune detection [8]. Among exosomal cargoes, miRNAs play a very crucial role in cancer progression. These miRNAs, known as “exomiRs”, play a functional role in PMN formation leading to cancer metastasis [9], immune evasion [10], macrophage polarization [11], drug resistance [12], and TME remodeling [13]. A widely studied exomiR –– miR-21, is often upregulated in various cancers such as breast, colon, and lung, and is associated with tumor growth, invasion, migration, and chemoresistance by regulating PI3K/Akt pathway [14]. Likewise, miR-221/222 are upregulated in many cancers such as liver, breast, and prostate, and are also associated with chemoresistance to Tamoxifen by inhibiting p27 and ERα production in breast cancer cells [15]. Another such exomiRs is miR-10b, which promotes metastasis in breast cancer by targeting a tumor suppressor gene, *HOX10* [16].

Despite many studies, a comprehensive pan-cancer analysis of exomiRs, their targets, and their potential functional roles in PMN formation and metastasis is missing. Here, we characterize exomiRs in the PMN across 7 cancer types, identifying high confidence exomiRs and their targets. We observed high GC content in exomiRs, and their targets were linked to pan-cancer as well as cancer-specific pathways. ExomiR targets were enriched for tumor suppressor genes, downregulated in PMN, and associated with survival. A support vector machine (SVM) learning model based on exomiR targets as features predicted therapy response in patient with high accuracy. Overall, we performed a systematic investigation of the role of exomiRs in cancer progression and metastasis, exploring their potential utility as therapeutic targets or biomarkers.

## 2. Methodology

### 2.1. Dataset Creation

ExomiR expression profiles were collected for 7 different cancer types ––breast (BRCA), colon (COAD), esophagus (ESCA), liver (LICH), kidney (KIRC), lung (LUAD) and pancreatic (PAAD). These profiles were collected either from cell lines or from tumors using either microarray or RNAseq technology (Table 1). We classified genome wide ∼2000 miRNAs as an exomiRs or non-exomiRs based on the following criteria. First, if the control data was provided, all miRNAs that were significantly (logFC >= 2 and adj-P <= 0.05) upregulated in the exosomes derived from cancer cell line or tumor versus the control were deemed as exomiR. Second, in the absence of control data, we considered the author-provided list of exosome-derived miRNAs as exomiRs.

**Table 1:**
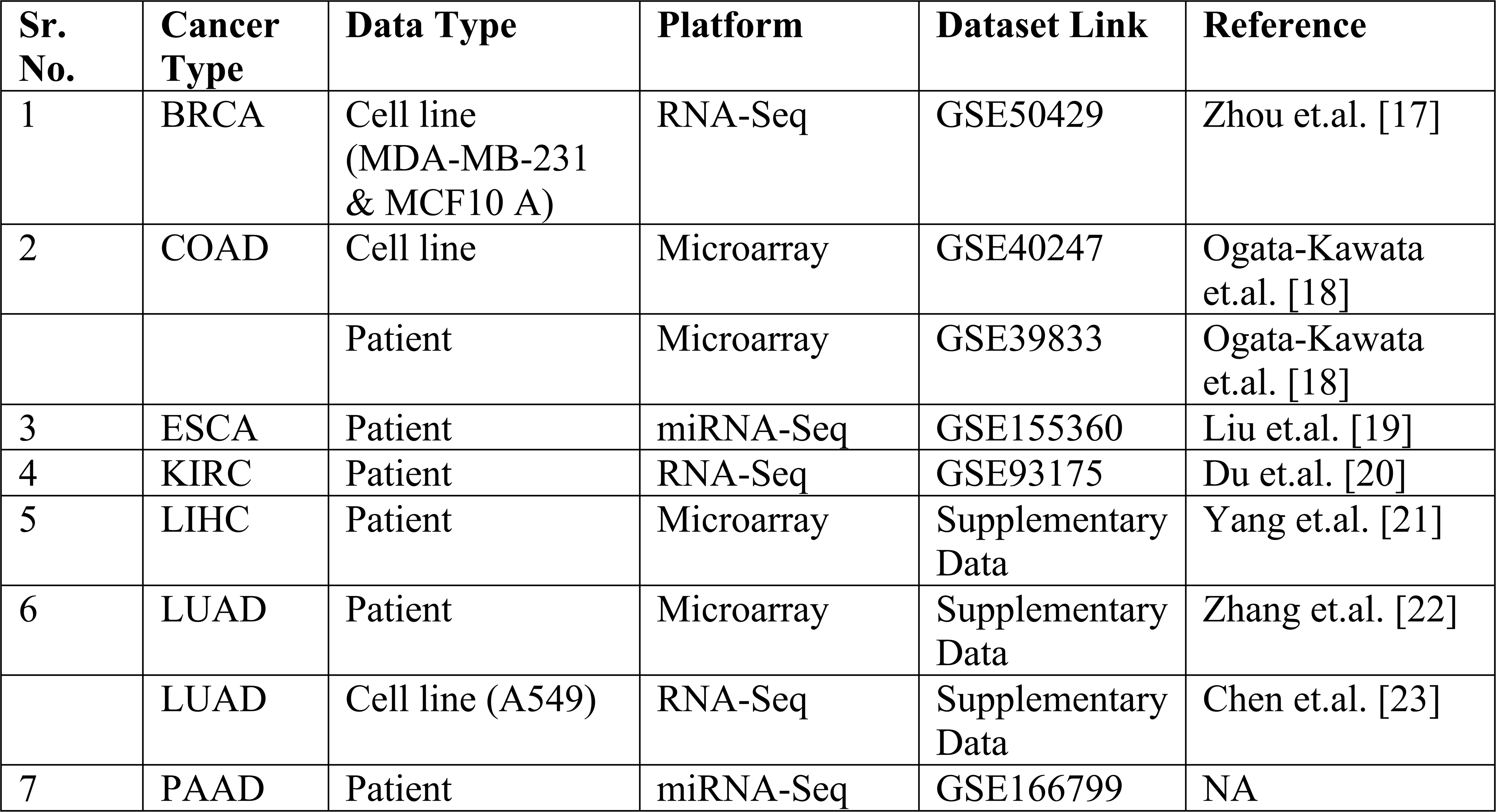
Data Description used in the current study. Here we enlist all the datasets which were analysed in this study across the cancer types.

### 2.2 De novo Motif Characterization from exomiRs

Using the miRBaseConverter package in R [24], we first converted the set of exomiRs found for each cancer type to their corresponding miRBase v22 identities to detect and compare context-specific exomiR motifs. Using the same program, we then obtained the sequences of the miRNAs that matched these miRBase v22 identifiers. We performed motif enrichment using MEME [25]. MEME tool was used for motif enrichment analysis using default RNA parameters that included a minimum motif width of 6 nucleotides, a maximum width of 50 nucleotides, and a 0-order background model. Pairwise motif comparisons were performed using TOMTOM [26] with default parameters, which include Pearson correlation for the motif comparison function, an E-value significance threshold of < 10, and complete scoring, to comprehend the similarity between the exomiRs motifs across our datasets (e.g., BRCA and LIHC).

### 2.3 Curating exomiRs and their putative targets

To further reduce potential false positives, we filtered the exomiRs based on GC content, as it has been reported, and as we observe, that exomiRs tend to be GC-rich [27]. We selected the exomiRs with GC content > 50%. Next, we characterized the experimentally ascertained targets of the exomiRs from the mirTarBase database [28]. Given the relative lack of specificity of miRNA targets, we further filtered the targets as follows. First, we characterized the significant enriched biological processes associated with those targets using ‘Enrichr’ tool [29] and next we computed the frequency with which each target belonged to the enriched processes and based on those frequencies, we selected the top 100 targets in each cancer type. These top 100 targets were used for all downstream analyses.

### 2.4 Gene Ontology analysis

ClusterProfiler 4.0 software [30] was used to analyze enriched biological processes (BP) associated with exomiR targets. We used top 100 targets as foreground and the default background. “Human” database was used as the background database, minimum and maximum gene size was set as 10 and 500 respectively to ensure specific terms. We used ‘simplify’ as function, ‘0.8’ as cutoff value to remove redundant terms and “Wang” as measure. Below we have provided the commands used to get enriched terms:

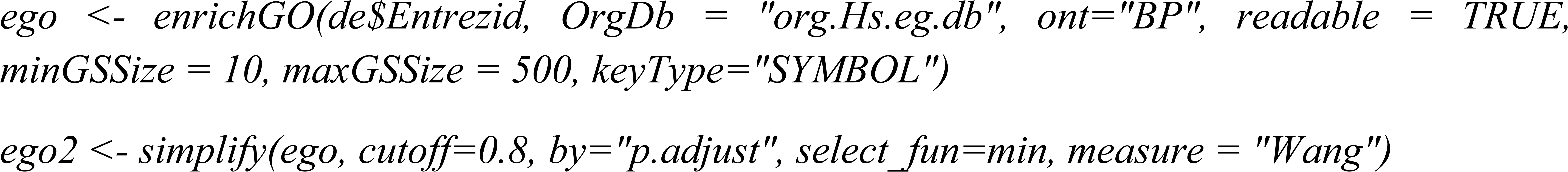

Next, we used the “Enrichr” website (https://maayanlab.cloud/Enrichr/) to perform KEGG analysis. List of exomiR targets were provided as an input and default parameters were used for the analysis.

### 2.5 Characterizing tumor suppressor genes properties of exomiR targets

To assess the enrichment of cancer specific tumor suppressor genes (TSG) in our exomiR target list, we extracted the list of cancer-specific TSGs from TSGene 2.0 database [31] TSGene 2.0 comprises information for 1217 human TSGs (1018 protein-coding and 99 non-coding genes) curated from more than 9000 articles. We were able to curate TSGs for 4 cancer types only which include BRCA, COAD, KIRC and LUAD as for the remaining 3 cancer types, data was not present in the TSGene 2.0 database. Next, we overlapped our top 100 exomiR targets with the TSGs in a cancer type-specific manner using Fisher’s Exact Test and computed Odd’s Ratio (OR) and statistical significance.

### 2.6 Expression pattern of exomiR targets in pre-metastatic niche (PMN)

We downloaded the gene expression data from the TCGA-TARGET project where TCGA and GTEx data have been processed uniformly and normalized collectively [32]. The tumor-adjacent morphologically normal tissues were taken as a proxy representative for pre-cancer or pre-metastatic niche (PMN) and the expression of exomiR targets were compared between the PMN and the healthy control samples from GTEx. We analyzed differential expression of our top 100 exomiR targets in PMN versus control using Log2(PMN/Control).

### 2.7 Survival analysis using gene expression of exomiR targets

We downloaded the gene expression and survival data from the TCGA database for respective cancer types. Cox proportional hazard model was used to compute Hazard Ratio (HR) using the gene expression as a feature using R package ‘survival’ [33]. First, we computed HR for all the top 100 exomiR targets and selected the one with negative HR and significant p-value (<0.05). This set was considered as “Observed”. Next, we performed the same analysis using all the coding and non-coding genes and termed it as ‘Expected’. Finally, we computed the Log (Observed fraction/Expected fraction) ratio to assess whether exomiR targets are more likely to be associated with better survival in comparison to random genes. We also performed Kaplan-Meier analysis [34] to highlight survival association with top exomiR targets with negative HRs in each cancer type.

### 2.8 Characterizing the role of exomiR targets in M2 macrophage differentiation

Macrophages are classified into 2 classes –– M1 and M2. M1 and M2 phenotype associated functions are classified into anti-tumor and pro-tumor respectively. Differentiation of macrophage into M1 or M2 phenotype depends on various signaling pathways. To elucidate the role of exomiR targets in pathways associated with differentiation of macrophage into M2 phenotype, we compiled list of such pathways from various studies [11,35,36] and assessed their overlap with our exomiR target lists.

### 2.9. Predicting therapy response based on exomiR targets

We compiled clinical datasets where for a given cancer type, chemotherapy treatment was administered and the information regarding the response to the given treatment is provided. We were able to get such data for 4 cancer types i.e. BRCA, COAD, ESCA and LIHC. For BRCA and COAD we obtained 2 such datasets whereas for ESCA and LIHC we obtained 1 dataset each (Table 2). Next, using python ‘scikit-learn’ package [37] we trained machine learning models, based on top 100 exomiR targets as features, to distinguish responder from non-responders in each clinical dataset. We used Support Vector Machine (SVM) and Random Forest (RF) machine learning approaches, as they have already been previously shown to perform well on such datasets [38]. We compared the performance of these models based on exomiR targets with other widely used signatures from a previous study [39] which include cancer-associated fibroblasts (CAFs), T cell exhaustion, immunotherapy targets, tumor associated macrophages (TAM) and tumor microenvironment (TME) signatures.

**Table 2:**
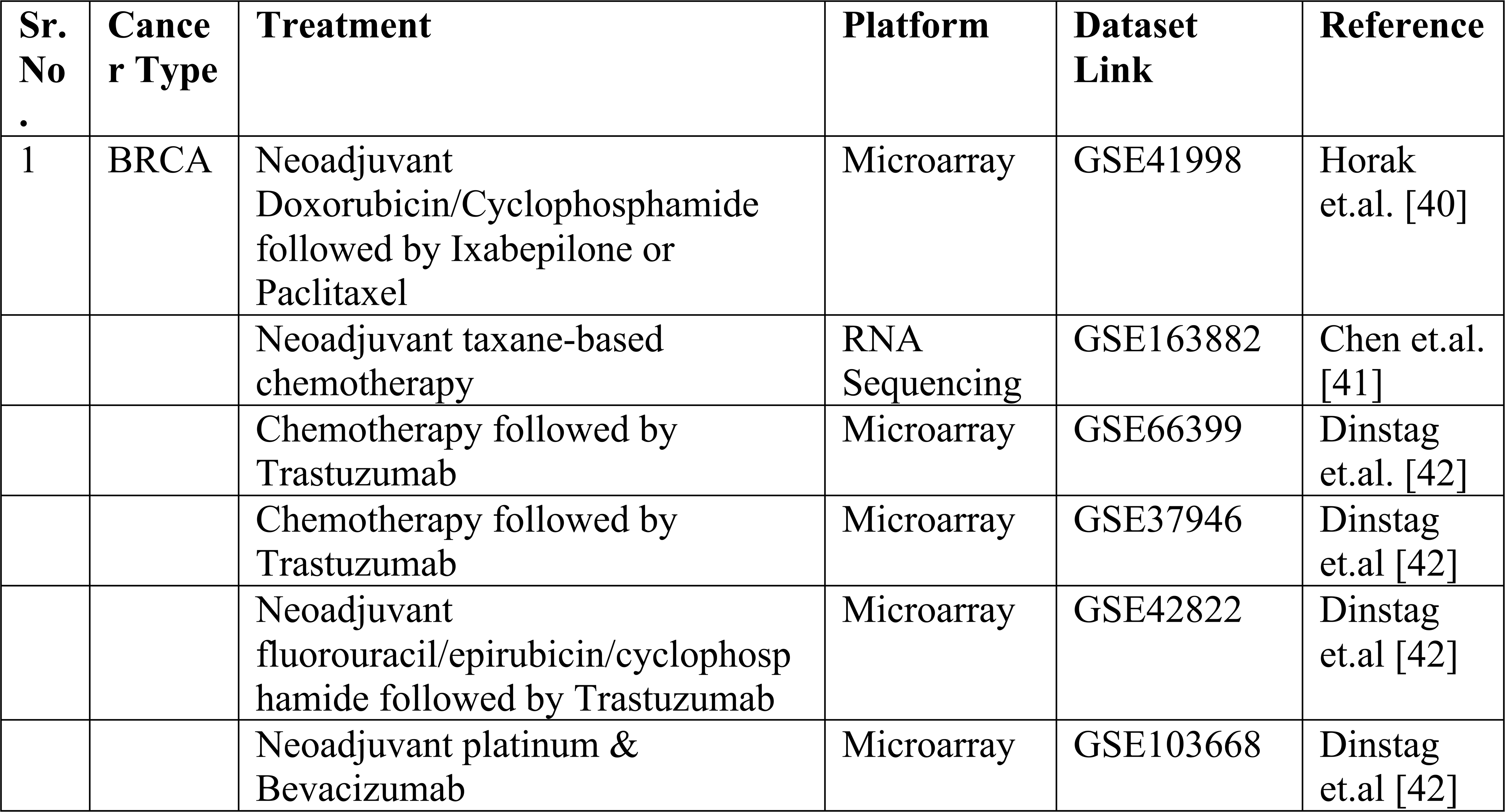
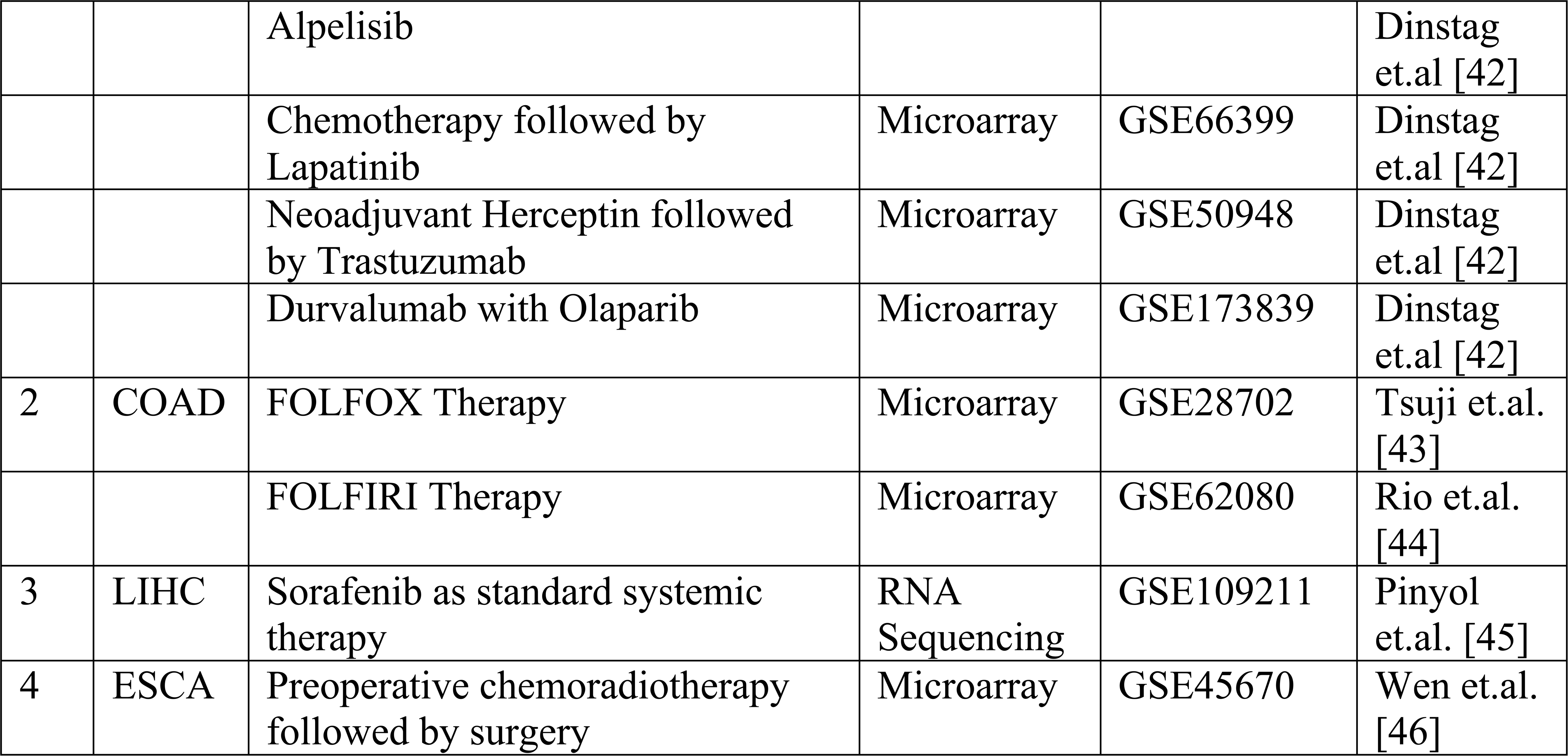
Data Descriptions Used for the Machine Learning Study. This table enlists all the datasets used during the responder/non-responder prediction across different cancer types.

## 3. Results

### 3.1 ExomiRs and their targets are rich in GC content and are enriched for specific sequence motifs

GC content is likely to be an important aspect of exomiRs’ role in cancer as high GC content is associated with the stability of the miRNAs and their target mRNAs, influencing the specificity and binding affinity of miRNAs to their targets [47,48]. High GC content exomiRs may thus have a more sustained regulatory effect on target genes in recipient cells [49]. Here we assessed the extent to which the exomiRs exhibit GC bias. As shown in Figure 1A, GC content of exomiRs is significantly higher than genome-wide miRNA background. Given this trend, to further narrow down our list of exomiRs, we retained the exomiRs with GC >50%. List of exomiRs along with their GC content (%) for individual cancer type is provided in Supplementary Table S1.

**Figure 1.**
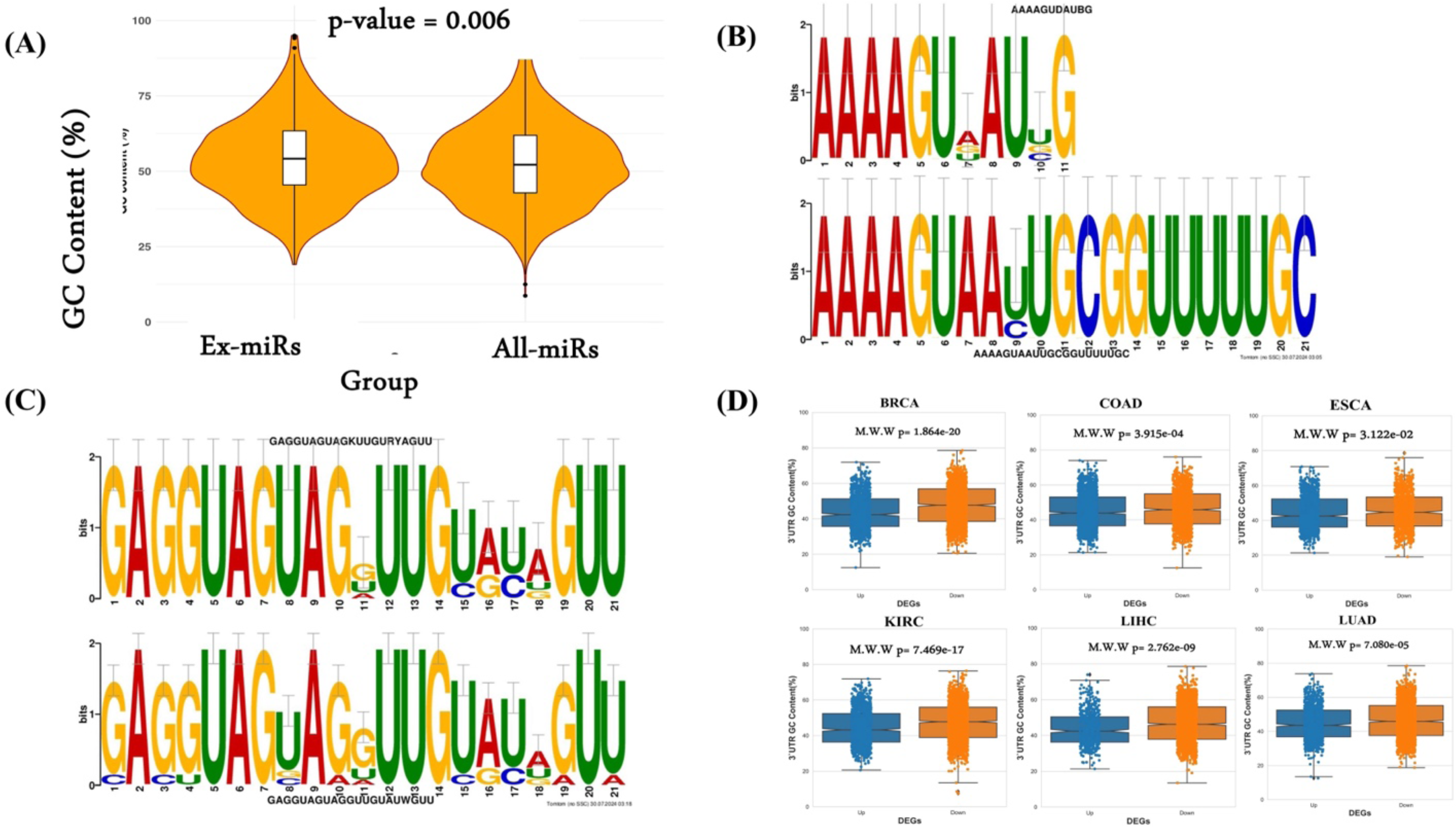
GC content and motif analysis. (A) GC content analysis of exomiRs characterized from various cancer types and complete mature human miRNAs. (B) TOMTOM alignment results for the paired motifs between characterized from LIHC-LUAD exomiRs and (C) KIRC – ESCC exomiRs. (D) GC content analysis of 3’ UTR region of downregulated genes in TCGA-NATs compared to GTEx with upregulated genes.

Beyond GC content, we also investigated sequence patterns enriched in exomiRs using MEME tool (Methods; Supplementary Table S2). We found that mononucleotide [UUU/AAA/GGG] and dinucleotide [CACA/AGAG] repeats were broadly enriched in exomiRs from various cancer types. ‘UG’ motifs were consistently identified in the basal portions of exomiRs, whereas motifs generated from esophageal cancer (ESCA) exhibited constant ‘UGU’ motifs in their apical regions. These apical ‘UGU’ and basal ‘UG’ motifs in pri-miRNAs are known to interact with the DROSHA and DGCR8 complexes [50]. ExomiR motifs were largely enriched for the ‘AGG’ motif across a variety of cancer types (except LIHC). However, LIHC exomiRs show enrichment for the ‘CCUC’ motif, consistent with the previous finding by Garcia-Martin et al., who also identified a similar motif (CC[C/U]C) in exomiRs from the AML12 cell line (alpha murine liver 12). In addition, we observe a frequent occurrence of the UUU (G/U) pattern in most of our dataset. One of the previous studies by Rolle et al.’s [51] has shown that miRNAs with UUU (G/U) motifs are associated with the neurotrophin signaling pathway, which is regulated through many intracellular signaling cascades, including the MAPK, PI3K, and PLC pathways, consistent with expected role of exomiRs in metastasis. Subsequently, we compared the enriched motifs using the TOMTOM tool. In general, we observed sequence similarity among the motifs characterized by exomiRs across cancer types. We have shown two such examples where sequence similarity can be seen among the motifs characterized by (i) LIHC and LUAD exomiRs (Figure 1B) and (ii) KIRC-ESCC exomiRs (Figure 1C). Complete comparison of enriched motifs identified in each cancer type with those in others is provided in Supplementary Table S3.

Assuming a role of exomiRs in shaping the tumor microenvironment, and reasoning that the genes downregulated in tumor-adjacent tissue may, in part, be targets of exomiRs, we computed the GC content (%) of the 3’ UTRs (the substrate for miR targeting) of the genes downregulated in TCGA-NATs compared to GTEx. As shown in Figure 1D, we observed significantly higher GC content in the 3’ UTR of the downregulated genes in all the cancer (PAAD does not have NAT data) with high significance. The high confidence exomiRs and their experimentally ascertained target genes were used in the downstream analyses.

### 3.2 ExomiR targets show shared and cancer specific gene ontology

Here, we investigated the extent to which various biological processes and pathways are enriched among the top 100 exomiR targets in a cancer type-specific manner (Supplementary Table S4). First, we characterized biological processes enriched among the targets using clusterProfiler 4.0. We observed shared as well as cancer-specific unique processes. Some key shared processes are epithelial cell proliferation, response to hypoxia, and positive regulation of miRNA metabolic processes. Previous studies have shown the role of these processes in cancer metastasis. For example, downregulation of TGF-β can alter the epithelial to mesenchymal transition (EMT) pathway and leads to cell growth, proliferation, and metastasis [52], promote oncogenic pathways such as MAPK/ERK and PI3K/AKT [53]. Likewise, various studies have shown the downregulation of components of hypoxic response promotes cancer metastasis, such as STAT3 induced EMT pathway [54], disrupting endothelial tight junctions, etc. [55].

Likewise, we also observed cancer-specific processes where the majority of them have been previously reported in various studies. For example, BRCA exomiR targets showed enrichment for apoptosis signaling and pathways related to the cell cycle [56–58]. COAD exhibited enrichment in processes linked to mitochondria [59,60]; ESCA displayed enrichment in processes related to peptidyl-tyrosine [61,62]; KIRC exomiR targets were associated with TGF-β signaling and kinase activity [63,64]; LIHC showed enrichment in processes related to smooth muscle cell proliferation [65,66]; LUAD was specifically associated with ossification and the Wnt signaling pathway [67,68]; PAAD exhibited enrichment in signaling cascades such as the MAPK cascade, and ERK1/ERK2 cascade [69,70]. Top 10 enriched processes for common and cancer specific ex-miR targets is shown in (Figure 2A-H) and complete list is provided in the Supplementary Table S5-S12.

**Figure 2.**
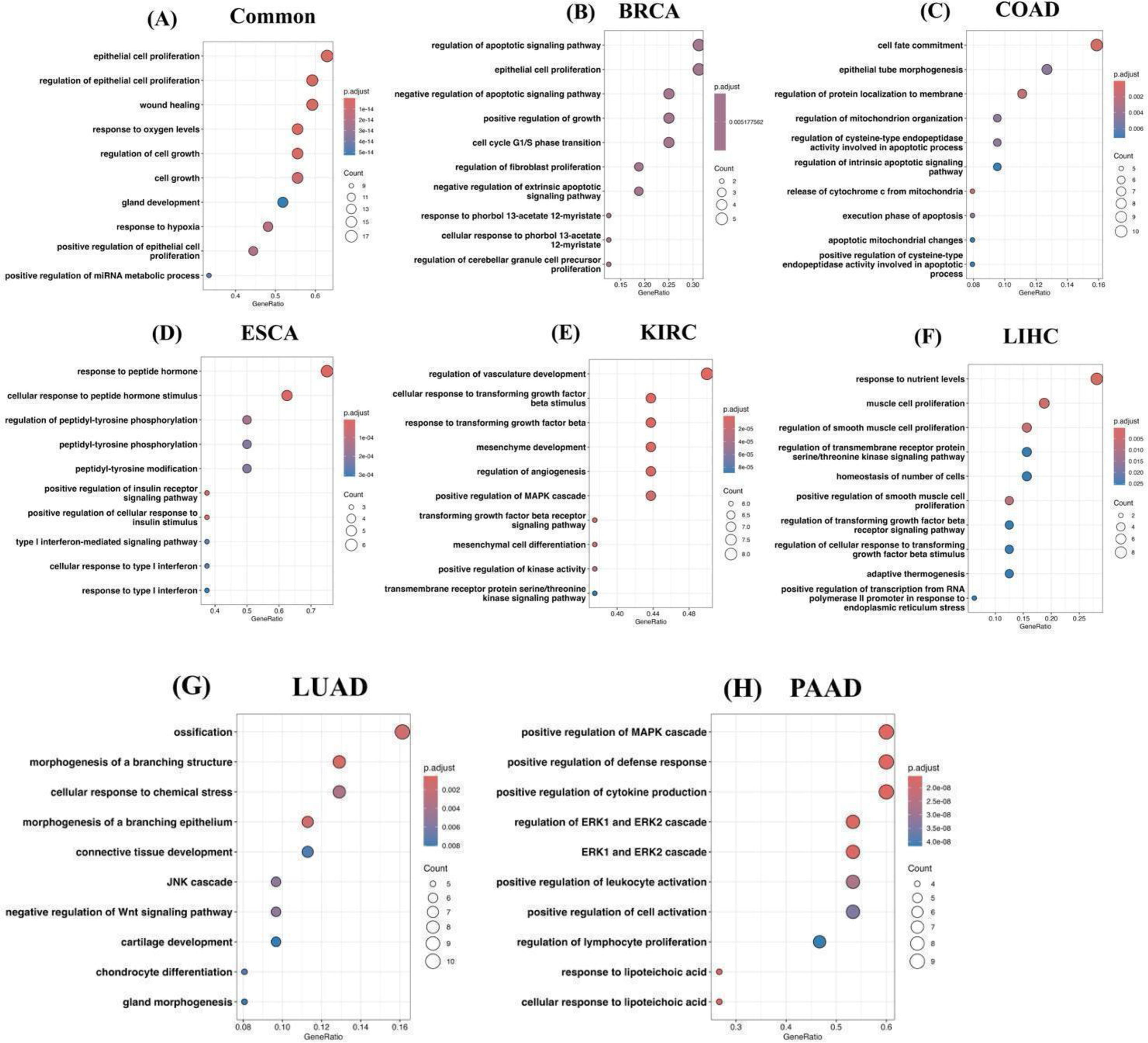
exomiR targets are enriched with essential biological processes. Top 10 enriched biological processes associated with common exomiR targets (A); unique exomiR targets in BRCA (B); unique exomiR targets in COAD (C); unique exomiR targets in ESCA (D); unique exomiR targets in KIRC (E); unique exomiR targets in LIHC (F); unique exomiR targets in LUAD (G); and unique exomiR targets in PAAD (H).

KEGG pathway enrichment analysis revealed significant enrichment of pathways such as “Pathways in cancer, MicroRNAs in cancer, HIF-1 signaling pathway, PI3K-Akt signaling pathway, Wnt signaling pathway, TGF-β signaling, etc. Role of these pathways in metastasis is previously established. For example, exomiRs miR-210, is upregulated in hypoxic condition and modulates the expression of various HIF-1 target genes involved in angiogenesis and EMT [71]. Likewise, exomiR miR-21 and mir-222 target *PTEN* which is a negative regulator of the PI3K-Akt pathway, leading to its downregulation and activation of the pathway which ultimately leads to cancer cell survival and proliferation [72]. Similarly, Wnt pathway is also regulated by exomiR such as miR-92a and miR-27a which targets Dickkopf-3 (DKK3), an inhibitor of Wnt signaling, leading to metastasis [73].

Top20 enriched KEGG pathways associated with common and unique genes among cancer types is provided in Supplementary Figure S1 whereas complete list of the common and cancer specific significant KEGG pathways is provided in Supplementary Table S13-S20.

### 3.3 ExomiR targets are enriched for tumor suppressor genes

Given the potential role of tumor exomiRs in promoting cancer, and the suppressive role of miRs, we assessed whether exomiR targets are enriched for Tumor Suppressor Genes (TSGs). We obtained the list of cancer-specific TSGs in four cancer types –– BRCA, COAD, KIRC and LUAD, and assessed their overlap with the exomiR targets using Fisher’s test. As shown in Figure 3, top 100 exomiR targets significantly overlap with the cancer-specific TSGs. Some of the key TSGs which were present among the targets in all four cancer types include *PTEN*, *SFRP1*, and *STAT3*. Previous studies have shown that downregulation of these genes by exomiRs are associated with cancer metastasis. For example, Zhang et.al. shows the role of *PTEN* loss by exomiRs primes brain metastasis outgrowth [74]; Wei et.al. shows ex-miR mir-221/222 contributes to tamoxifen resistance by targeting *SFRP1* in ER-positive breast cells thereby promoting tumor progression [15]; and Wang et.al. shows the role of *STAT3* downregulation in cancer progression [75]. Some other key TSGs overlapping exomiR targets include *SIRT1, SMAD2, FOXO2, TGF-β*, etc.

**Figure 3.**
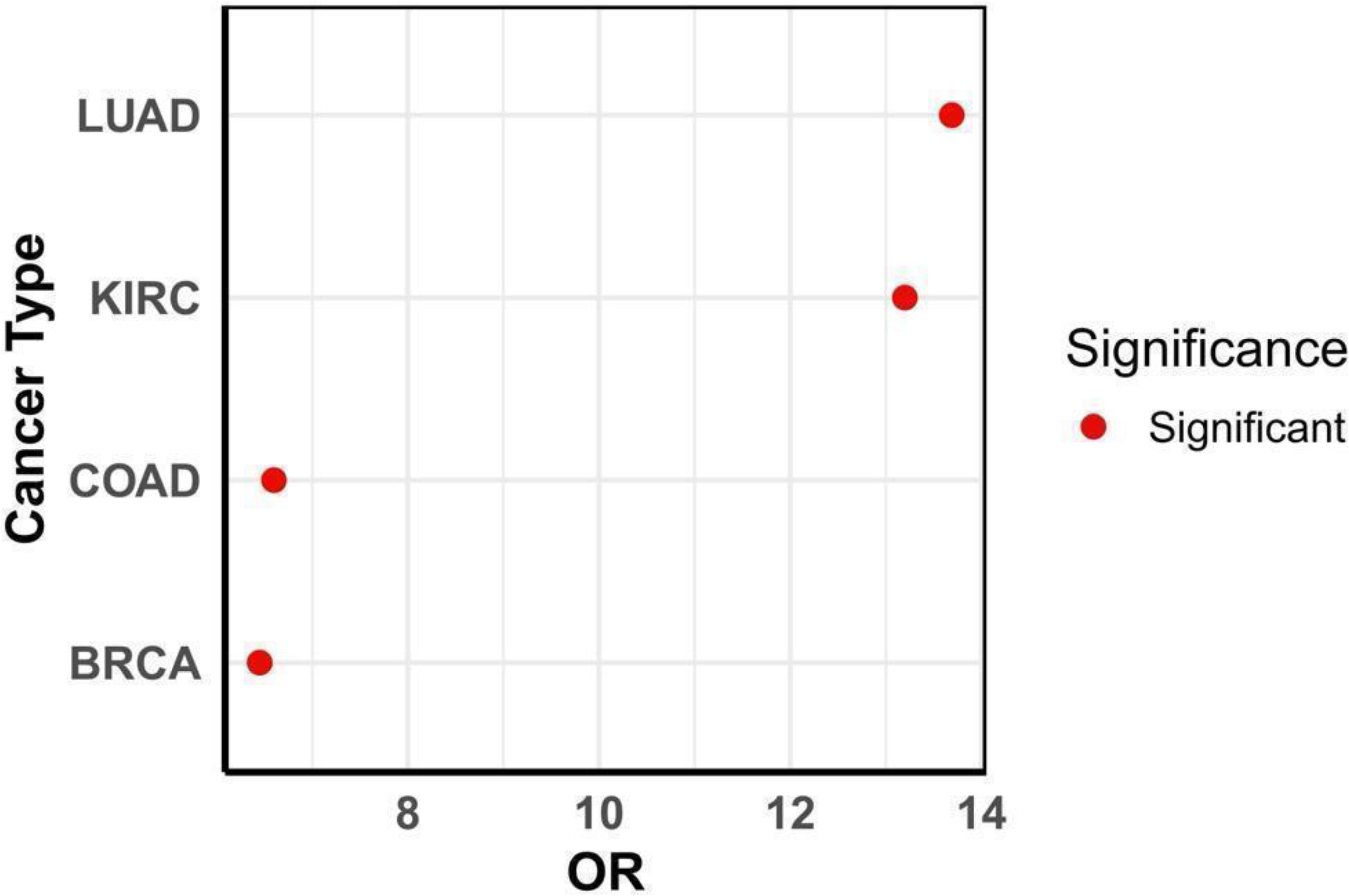
exomiR targets are enriched for Tumor Suppressor Genes. Fisher’s exact test Odd’s ratio shows statistically significant enrichment of TSG among exomiR targets in various cancer types.

### 3.4 ExomiR targets are downregulated in the pre-metastatic niche compared to healthy donors

Role of cancer derived exomiRs in PMN formation and remodeling is well established [76,77]. Here, we analyzed a dataset where PMN formation was observed [78]. We observed the downregulation of exomiR targets in PMN observed in lung tissue compared to healthy regions for all the cancer types except LUAD as represented in the violin plot (Figure 4). Log2(PMN/Control) gene expression data used is provided in Supplementary Table S21. This analysis strengthens our hypothesis that cancer derived exomiRs bind to their target genes in distal organs to contribute toward PMN formation, prior to metastatic colonization.

**Figure 4.**
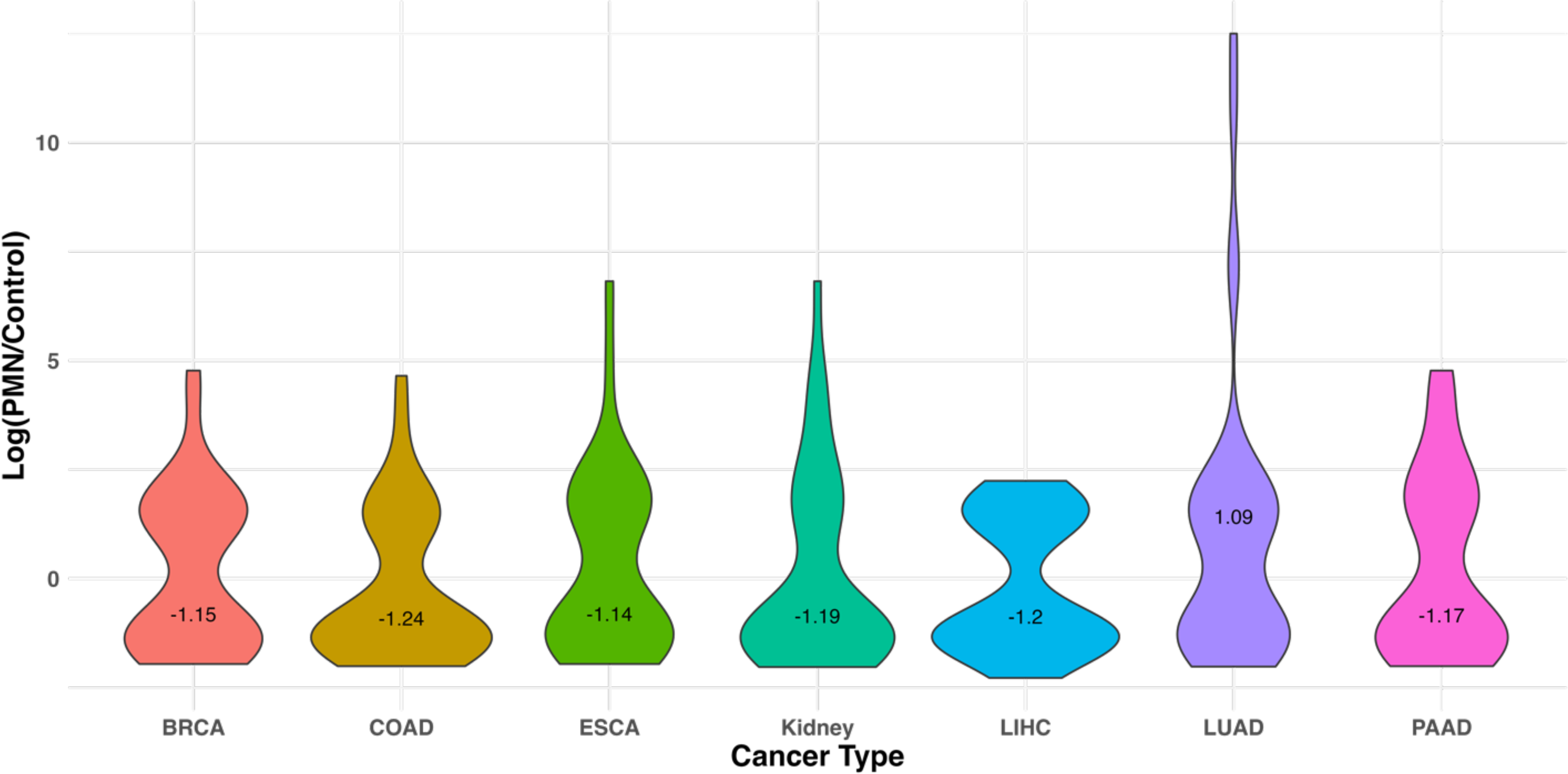
exomiR targets are downregulated in the pre-metastatic niche compared to normal. Violin plot showing median Log fold change of gene expression in PMN which formed in lung tissue, compared to normal in the mouse data with rhabdosarcoma. Median Log2FC value is printed on the plot.

### 3.5 ExomiR targets are associated with survival in some cancer types

Given that our analysis above shows exomiR targets to be enriched for TSGs, here, we assessed whether exomiR target expression is predictive of better patient survival using the TCGA cohorts. We computed Hazard Ratio (HR) for all the top 100 ex-miR targets in each cancer type using Cox proportional hazard model where gene expression of the gene was used as an input feature, controlling for age, sex, and tumor purity. Next, we noted the fraction of genes with significant (p

< 0.05) negative HR (i.e., their expression is associated with better prognosis), listed in Table 3; complete data for all the 100 exomiR targets in each cancer type is provided in Supplementary Table S22-S28. We compared the fraction of exomiR targets (Observed) with significant negative HR with the same for the genome-wide background (Expected) via log(Observed/Expected). As shown in Figure 5A we observed high enrichment of Observed targets compared to Expected targets associated with better survival outcome in all the cancer types except PAAD.

**Figure 5A:**
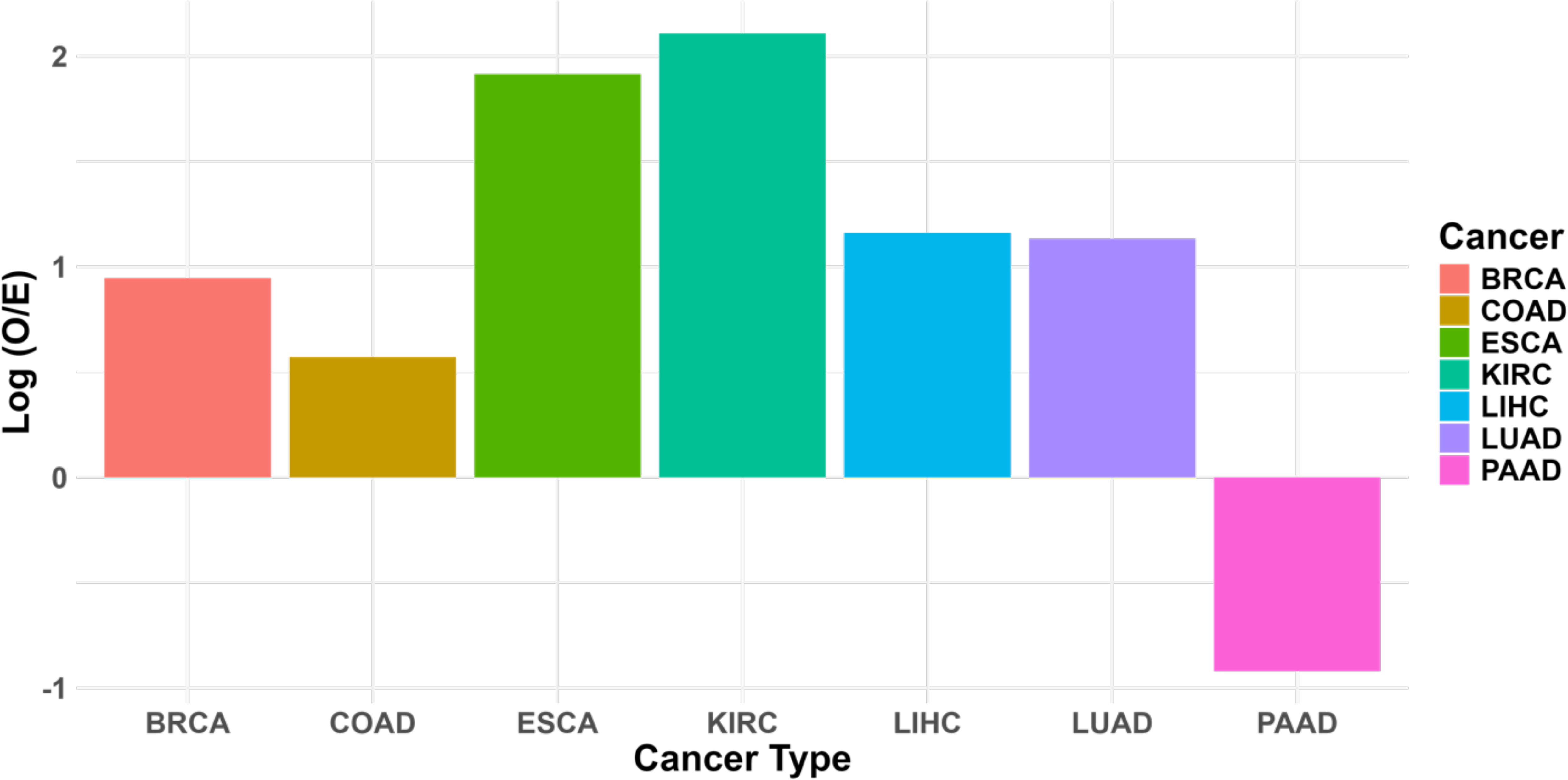
exomiR targets are associated with overall survival. Cox proportional hazard model shows that exomiR targets are more statistically significantly enriched for survival association compared to in general expectation in each cancer type. Log2FC (Observed/Expected) was computed where ‘Observed’ represents percentage of exomiR targets with negative HRs and p-value < 0.05, and ‘Expected’ represents percentage of any target (coding & non-coding) with negative HRs and p-value < 0.05.

**Table 3.**
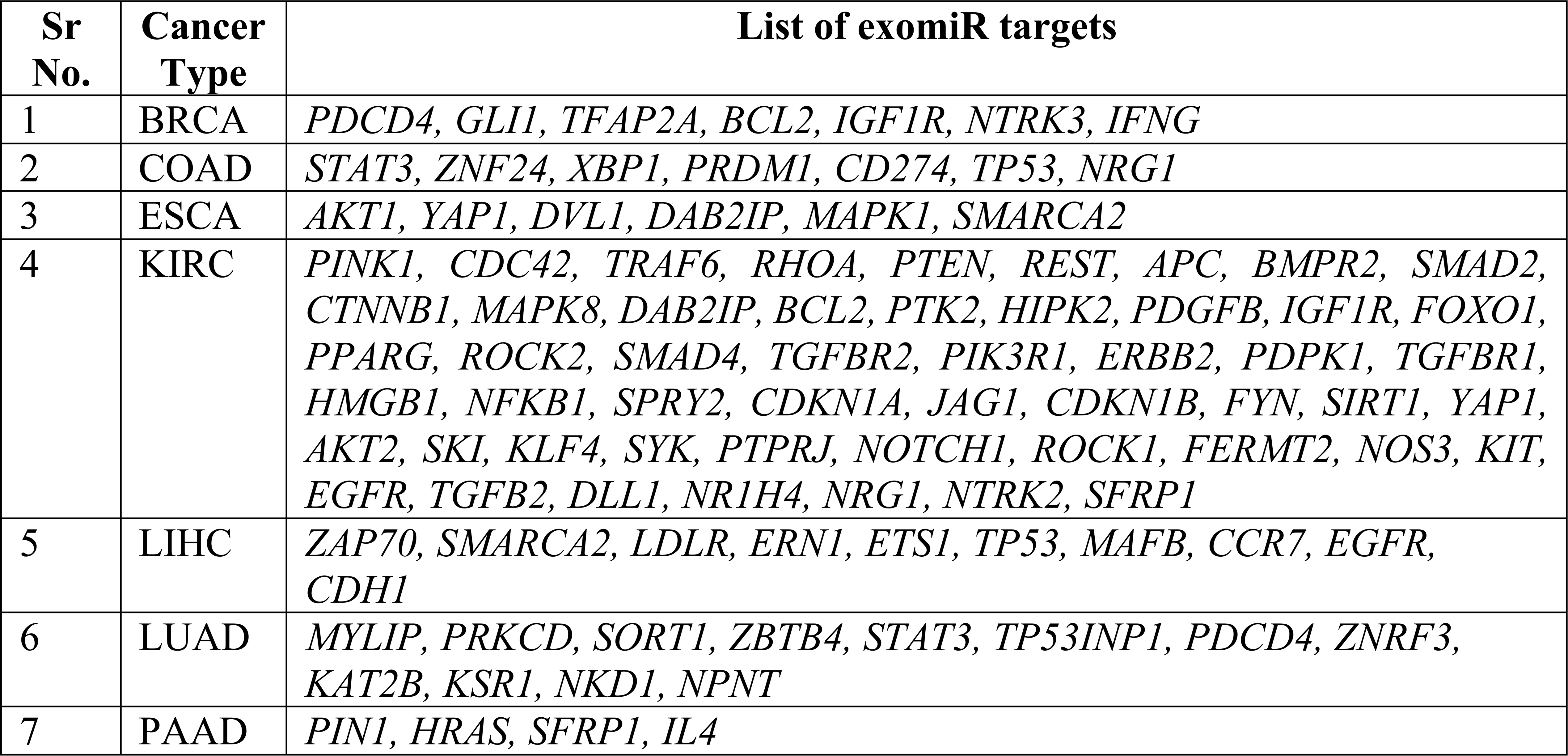
List of ex-miRs targets significantly associated with survival association. Here we provide list of all the exmiRs targets across various cancer types which shows negative Hazard Ratio and p-value (<0.05).

We further obtained the top 10 exomiR target genes with significant negative HR based on p-values, and for each gene, we partitioned the corresponding TCGA cohort in to ‘High’ and ‘Low’ (Top50% and Bottom 50%) categories based on the median gene’s expression and compared the differences in their survival using Kaplan-Meyer survival curves and estimated the log-rank p-values. We observed that KIRC performed best among all the 7 cancer types with all 10 genes showing significant association with survival, followed by PAAD with 75% of the genes showing significance (3/4) and BRCA with 57% genes (4/7) showing significance. In the case of LUAD, we observed 4/10 genes with significant p-value. In case of COAD and ESCA only 1 gene shows the significant p-value. One example from each cancer type is provided in Figure 5B and results for remaining genes are provided in Supplementary Figure S2-S8.

**Figure 5B.**
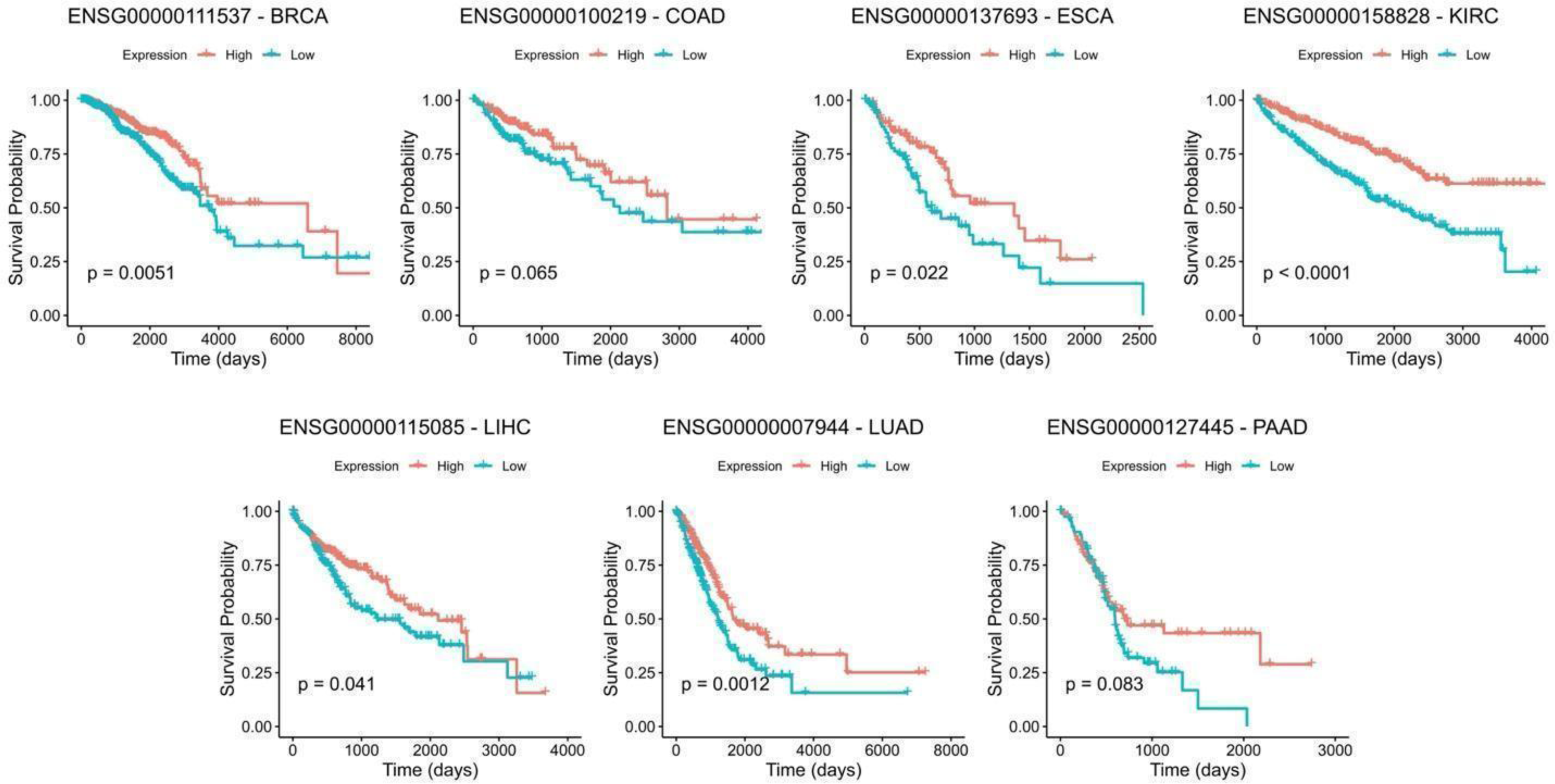
Kaplan Meier Survival Analysis. Kaplan Meier curve of the top genes selected from cox-regression analysis across cancer types. For each gene, patients were stratified into high and low category based on median expression and p-values were estimated using log-rank test.

### 3.6 ExomiR targets are associated with M2 macrophage polarization

Macrophages are diverse and adaptable immune cells crucial in combating pathogenic microorganisms and tumor cells. Depending on the stimuli they receive, macrophages can polarize into the M1 phenotype, which exhibits pro-inflammatory properties and inhibit tumor growth, or the M2 phenotype, which displays anti-inflammatory characteristics and promote angiogenesis [11]. Balance of macrophage polarization is crucial for disease progression and there are several factors associated with the polarization, including epigenetic modifications [79,80], and key signaling pathways such as AKT [81], Notch [82], TGF-β/Smad [83], JAK-STAT [84].

Here we assessed whether exomiR targets play a role in macrophage polarization. We found that our exomiR targets capture key signaling pathways whose downregulation leads to M2 macrophage polarization, discussed next.

Xun et. al. has shown that in breast cancer, downregulation of *KDM6B* by miR-138-5p switches M1 to M2 macrophage [85]. Interestingly, we observed the *KDM6B* among the exomiR targets specifically in BRCA. Haydar et.al have shown that downregulation of *STAT1* and *NFKB* by azithromycin leads to M2 macrophage polarization [35]. In our analysis, we recapitulated *STAT1* in ESCA and *NFKB1* in BRCA, ESCA, KIRC and LICH. Likewise, we *PTEN* was among the top exomiR targets in all cancer types except LICH; downregulation of *PTEN* leads to PI3K/AKT signaling pathway which leads to M2 macrophage polarization, which in turn promotes metastasis by secreting *CXCL13* [86], promotes angiogenesis by secreting *VEGFA* and *MMP9* [87]. Likewise, we were also able to recapitulate exomiRs associated with M2 macrophage polarization under hypoxic conditions which includes miRNA-940, miRNA-21-3p, miRNA-125b, etc. as observed by Li et.al. [88]. Although these miRNAs were obtained from epithelial cells in ovarian cancer, it would be interesting to explore their role in cancer types we analyzed in our study.

We also identified members of transcription factors (TFs) families involved in macrophage polarization. For example, *KLF6* is involved in M2 macrophage polarization in atherosclerosis [89], however, we observed another member, viz, *KLF4* in multiple cancer types among exomiR targets; however, *KLF4* has not been directly implicated in M2 polarization and warrants further exploration. Similarly, the role of transcription factor *IRF5* in M2 macrophage polarization is reported in the case of spinal cord injury [90], whereas we observed IRF4 among the exomiR targets in COAD. *FoxO1* is another TF whose role has been widely studied in M2 macrophage polarization [36]. In our analysis, not only we found *FoxO1* among exomiR targets, but also another closely related protein *FoxO3*.

Overall, our exomiR targets recapitulated previously reported targets associated with M2 macrophage polarization in various diseases (including cancer) and point to involvement of closely related family members in the process. Table 4 lists some key miRNAs, and their targets associated with M2 macrophage polarization revealed by our analysis.

**Table 4.**
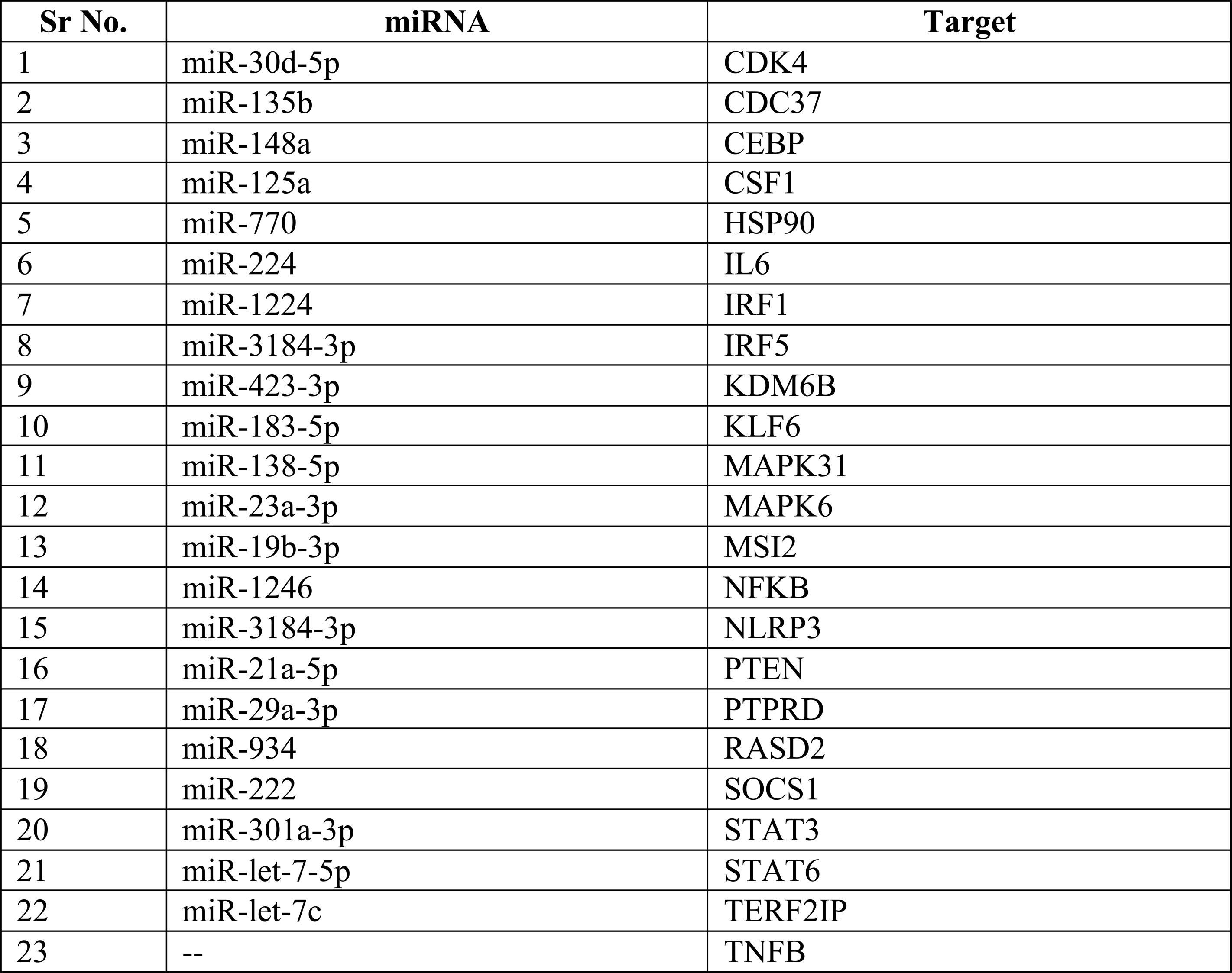
List of exomiRs and targets associated with M2 macrophage polarization. In this analysis, we provided all the exomiRs and their associated targets recapitulated in our study which when gets downregulated leads to M2 macrophage polarization phenotype.

### 3.7. ExomiR targets predict therapy response

Characterizing transcriptional signatures that can classify responder and non-responders to a given treatment to cancer patients is critical in clinical management. Table 5 lists the signatures used in the current study for comparison with exomiRs signatures. These signatures are (i) conventional Immune checkpoint inhibitors (ICI) such as ICI targets such as PD1, PD-L1 or cytotoxic T-lymphocyte antigen 4 (CTLA4) and (ii) tumor microenvironment associated markers such as CD8 T cell, T-cell exhaustion, cancer associated fibroblast (CAF), tumor associated macrophages (TAM) and Tumor microenvironment (TME) markers. These markers are well established and have been used in multiple studies to predict ICI treatment response in patients [39,91,92].

**Table 5:**
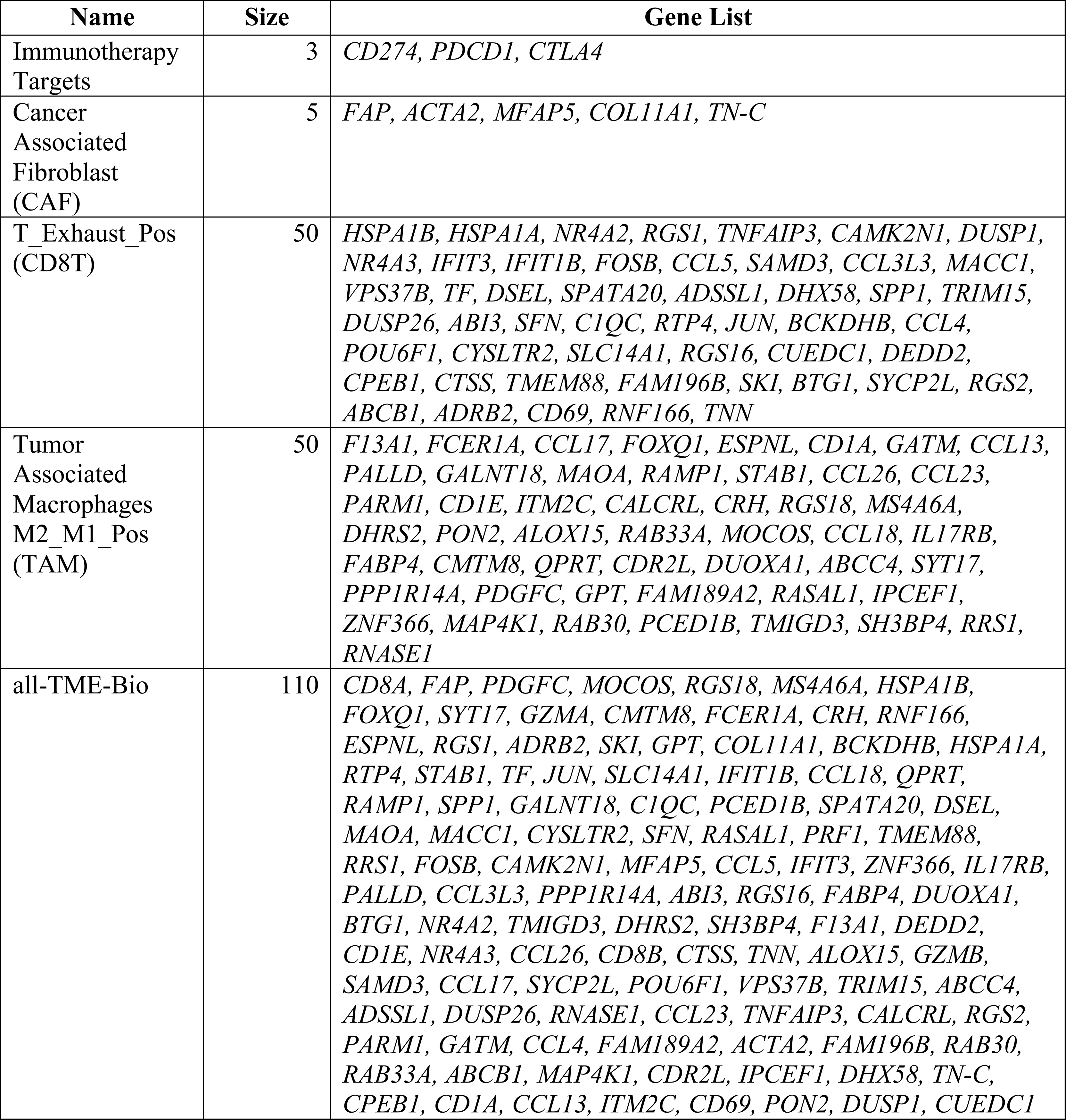
Gene Signatures used in the therapy response prediction. The Table enlist genes associated with each marker, used to compare with exomiR targets in predicting responder and non-responder to a treatment.

Here we evaluated the relative potential of exomiR targets to serve as gene signatures to discriminate responders from non-responders to ICIs. We collected several datasets from the literature where for a given cancer type, pre-treatment transcriptomic profiles of responder/non-responder to a given chemotherapy is provided. In the case of BRCA and COAD, there were two such datasets and for ESCA and LIHC there were one dataset each. We developed Support Vector Machine (SVM) models to discriminate responder/non-responders to given chemotherapy using the gene expression as a feature. Specifically, for BRCA, we procured eight additional datasets from the ENLIGHT study.

For BRCA, we trained the model on GSE41998 dataset and tested it on independent GSE163882 and the eight additional datasets from the ENLIGHT study. We observed that exomiRs targets performed favourably in a majority of cases (5 of the 9 datasets) whereas other signatures performed better on the remaining 4 datasets. In Figure 6A, we have shown the result for GSE163882, where our model trained using exomiR targets outperformed other gene signatures, with an AUROC of 0.72, followed by TME gene signature with AUROC of 0.67. Results on remaining BRCA datasets are shown in Supplementary Table S29. In the case of LIHC where we had only one dataset (GSE109211), we performed a 5-fold cross-validation. We observed that exomiR targets and CD8+ T cell signatures performed equally well with AUROC of 0.96. For COAD, we had two datasets (GSE28702 & GSE62080), however, both have very few samples. Hence, we combined these two datasets and performed 5-fold cross-validation. In this case our exomiR targets achieved 2^nd^ best performance with AUROC of 0.75 whereas CD8+ T cell signature performed best with AUROC of 0.85. Lastly, in the case of ESCA, we performed 5-fold cross validation on GSE45670. Again, exomiR targets performed best among all signatures with an AUROC of 0.83 followed by CD8+ T cells with AUROC of 0.78. The AUROC plot is shown in Figure 6(A-D) for the 4 cancer types and Supplementary Table S29 provides the information of the parameters of the SVM model.

**Figure 6.**
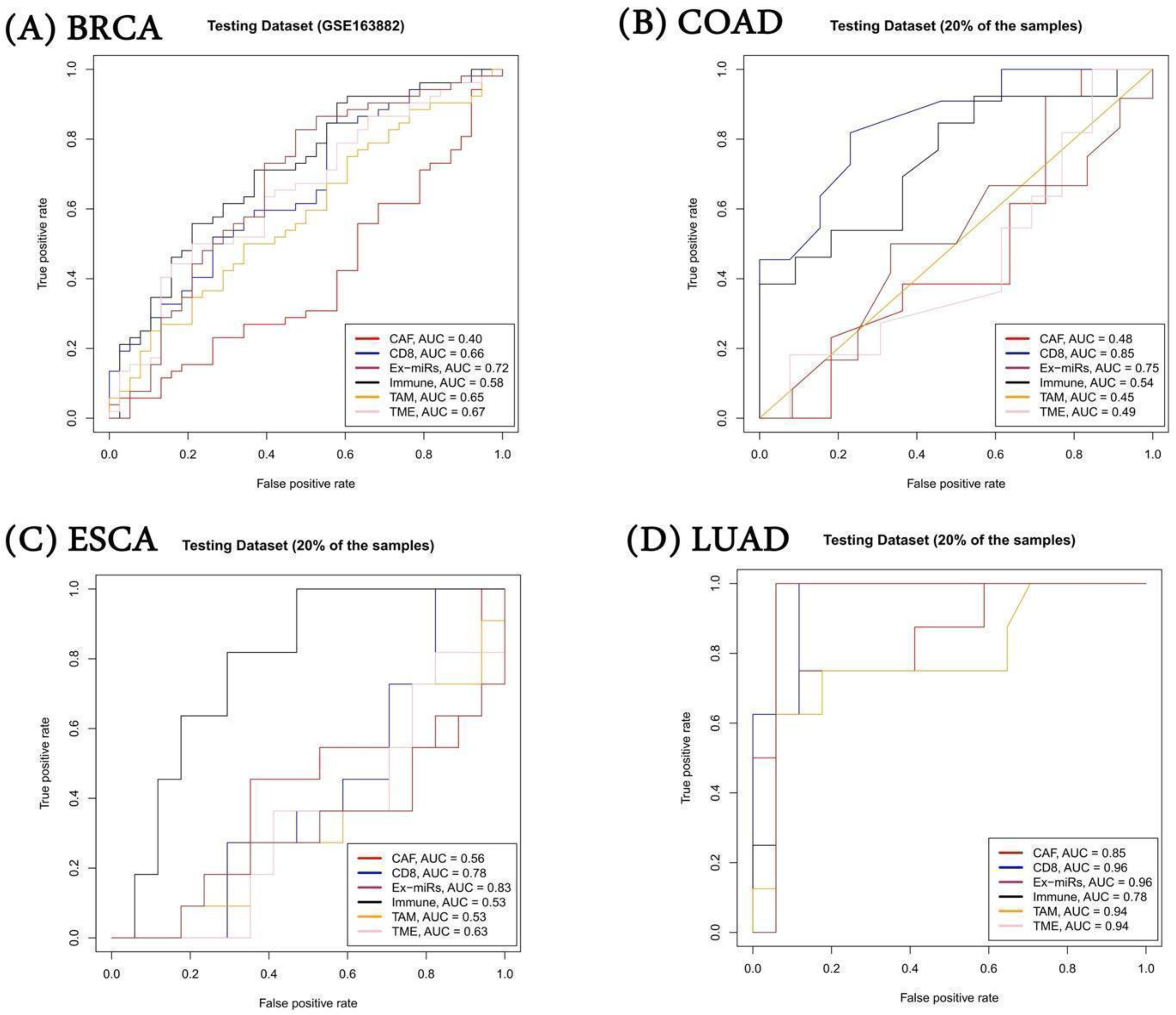
Performance of SVM model on independent dataset. Average gene expression of various gene signatures was used as a feature to build SVM models. exomiR targets were able to discriminate responder and non-responder significantly with high AUROC compared to other signatures in all cancer types except COAD where CD8+ signature shows highest AUROC followed by exomiR targets.

Overall, the above results shows that exomiR targets could be used as an alternative gene signature for classifying responder/non-responders treated with chemotherapy.

## Discussion

Intercellular communication is critical in homeostasis, and by extension, in disease. Exosomes, a membrane bound extracellular vesicles, are secreted by almost every cell, and carrying a variety of cargo, including miRNAs, they represent one of the primary modes of intercellular communication mediating numerous functions [93,94]. For example, downregulation of miR-122-5p, miR-590-5p in serum exosomal level has been seen in gastric cancer patients compared to healthy controls [95,96]. Likewise, the role of miR-15a secreted by mesenchymal cells has been shown to slow down hepatocellular carcinoma progression by downregulating *SALL4* [97].

Recently, there has been an exponential rise in the reports assessing the role of exosomal miRNAs in cancer [55]. The first report of miRNA in cancer was reported by Calin et al. in B-cell chronic lymphocytic leukemia (B-CLL) where they observed frequent deletion and downregulation of miR-15a/miR-16-1 which have tumor suppressor role [98]. miRNA-21 and miRNA-146a have immunosuppressive properties and can act on cervical cancer associated T-cells to promote oncogenesis [99]. ExomiRs also promote metastasis by aiding the formation of pre-metastatic niche (PMN) in distant organs. For example, exosomal miR-105 secreted by breast cancer cells is taken up by endothelial cells in distant organs, targeting and downregulating tight junction protein ZO-1. This leads to increased vascular permeability which allows cancer cells to metastasize in distant tissues particularly in lung and liver [17]. On the other hand, exomiRs present a therapeutic opportunity as well. For example, Hamideh et al. show that miRNA-21-sponge packed in exosomes has the potential to treat brain cancer [100]. Lastly, exomiRs have been found to be one of the factors associated with the developing resistance to various treatments against cancer. For example, Rodriguez-Martinez et.al showed that exosomal miRNA-222 downregulates PTEN pathway and leads to cell cycle arrest in breast cancer cells, conferring resistance to drug-sensitive cells [58].

With the increasing appreciation of the exosomes’ role in cancer, isolation and quantification of exosomes have become a crucial initiative in both basic research and clinical applications. Reliable and efficient isolation of exosomes from various biofluids (blood, urine, cerebrospinal fluids, etc.) is critical as next-generation biomarkers in various diseases [101]. Despite such technological advancements in exosome isolation, purification and sequencing of cargoes, the mechanism of how exosomal miRNAs regulate metastasis is not clear. One of the major issues is the lack of target specificity as single miRNA can target multiple targets or a single target can be targeted by multiple miRNAs, making miRNA-mRNA interactions complex to understand. Furthermore, the natural inter– and intra-tumor heterogeneity is likely to extend to exomiRs as well and thus, a more informative analysis of exomiRs’ role in metastasis would require a paired datasets of exomiRs from primary tumors and the metastatic microenvironment, which is currently not available.

In this comprehensive analysis of exomiRs from cancer cell lines and patient tumors for 7 cancer types, we observed common as well as cancer specific biological processes potentially regulated by the exomiRs. ExomiRs targets were found to be enriched for tumor suppressor genes and exhibited gene expression downregulation in pre-metastatic niche in single cell RNA-seq data. Survival analysis revealed significant enrichment of exomiRs targets in the genes associated with overall survival. In addition, exomiRs targets were found to be associated with M2 macrophage differentiation and overall better predictor of responder and non-responder to a given therapy compared to other markers.

Other than miRNAs, diverse range of molecules are packaged in exosomes, including long non-coding RNAs (lncRNAs), circular RNAs (circRNAs), proteins, lipids, metabolites, cytokines and chemokines. Like miRs, these other types of molecules are also associated with a wide range of functions. Isolation and characterization of those other information modalities, as the technology and data become available, will shed further light on the role of exosomes in oncogenesis, metastasis, and therapy resistance.

## Data and Code Availability

This paper analyzes existing, publicly available data. These accession numbers for the datasets are listed in the key resources table.

Any additional information required to reanalyze the data reported in this paper is available from the lead contact upon request.

Codes used for various analyses during the study is provided at our GitHub link https://github.com/agrawalpiyush-srm/Exosomal_miRNA/.

## Author contributions

PA and GO collected and processed the dataset. PA and GO perform the analysis. AS perform the survival analysis. VG process the single cell dataset. PA, GO and SH wrote the manuscript. PA and SH conceive the idea.

## Supporting information

All Supplementary Figures

All Supplementary Tables

## Acknowledgement

This research was supported by the Intramural Research Program of the NIH, USA and SRM Institute of Science & Technology, Kattankulathur, India.

## Declaration of generative AI and AI-assisted technologies in the writing process

During the preparation of this work the author do not used any AI or AI-assisted technologies to improve language and readability.

