## Supplementary material for "Characterizing the role of exosomal miRNAs in metastasis": All Supplementary Figures

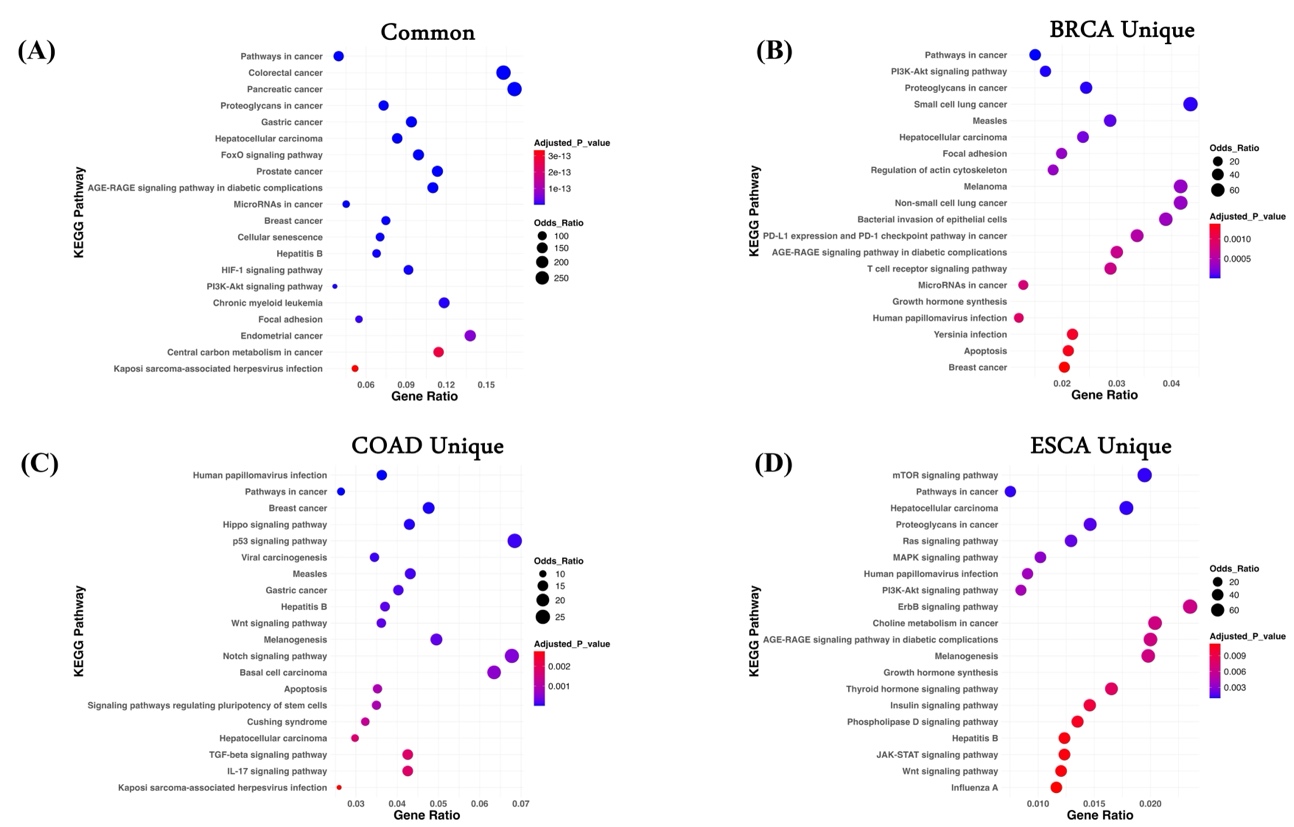

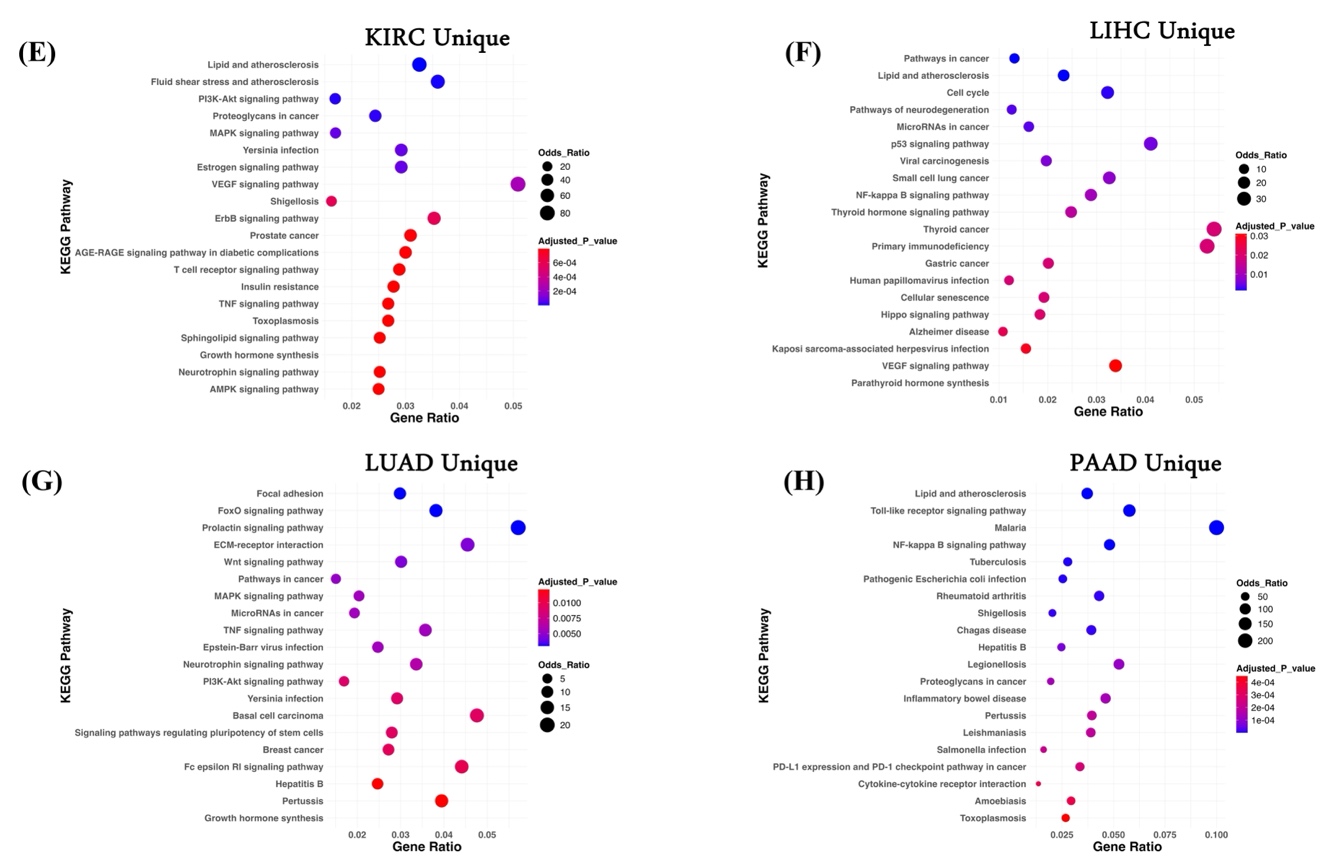

**Supplementary Figure 1. KEGG Pathway Analysis.** Top20 enriched KEGG pathways associated with common genes (A) among cancer types; (B) BRCA unique genes; (C) COAD unique genes; (D) ESCA unique genes; (E) KIRC unique genes; (F) LIHC unique genes; (G) LUAD unique genes; and (H) PAAD unique genes.

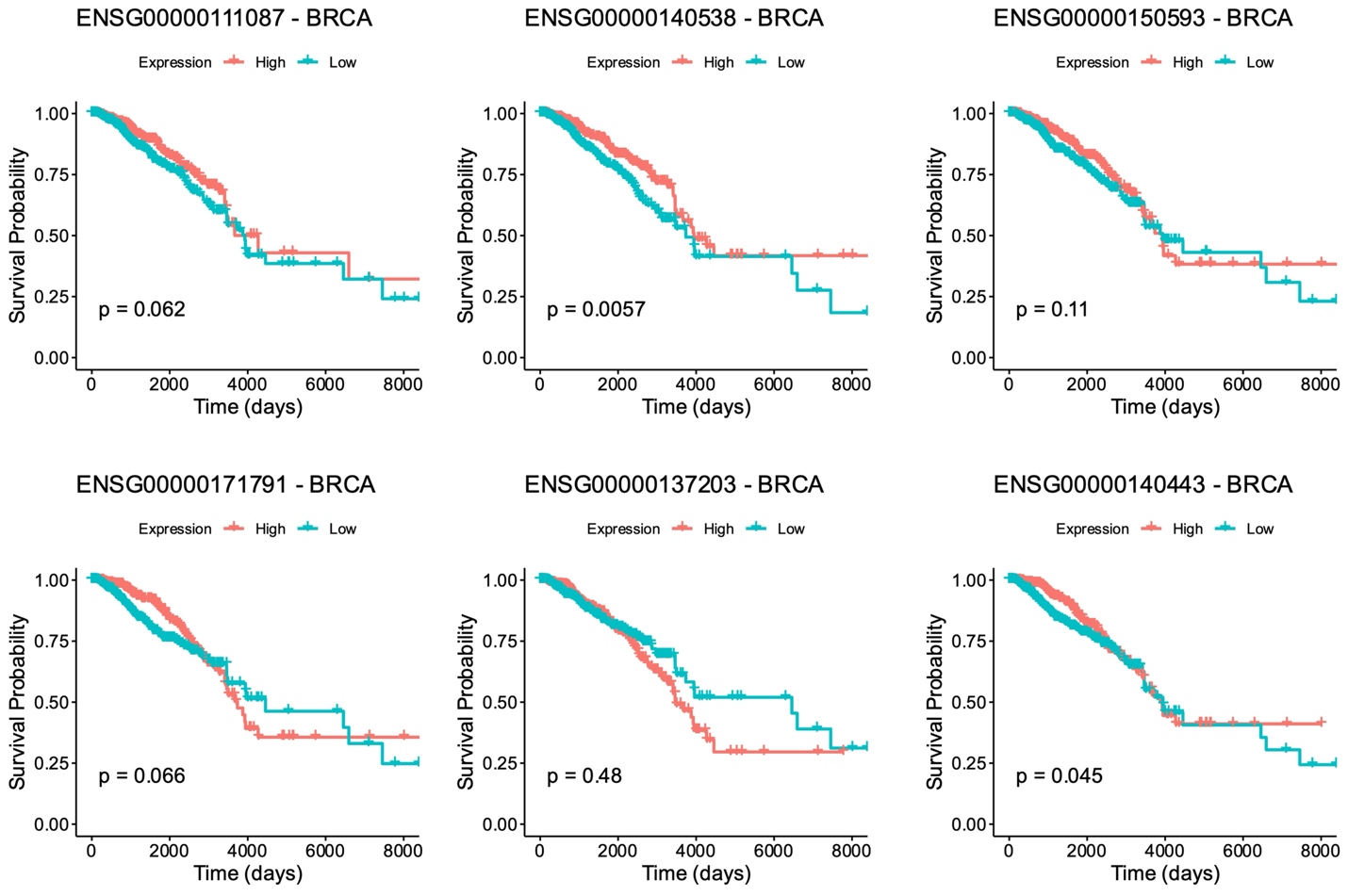

**Supplementary Figure S2.** Kaplan Meier curve of the top genes selected from cox-regression analysis in BRCA. For each gene, patients were stratified into high and low category based on median expression and p-values were estimated using log-rank test.

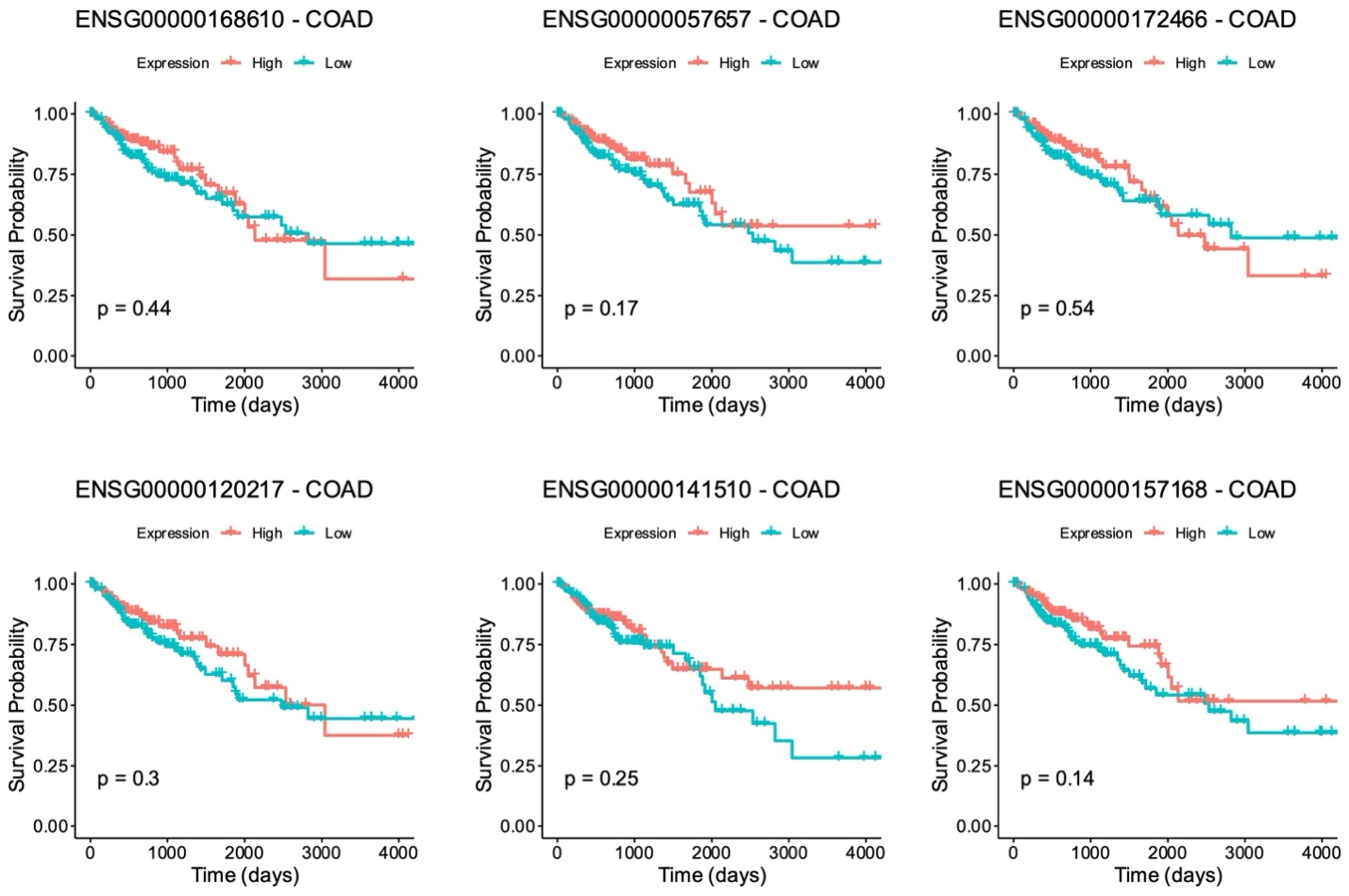

**Supplementary Figure S3.** Kaplan Meier curve of the top genes selected from cox-regression analysis in COAD. For each gene, patients were stratified into high and low category based on median expression and p-values were estimated using log-rank test.

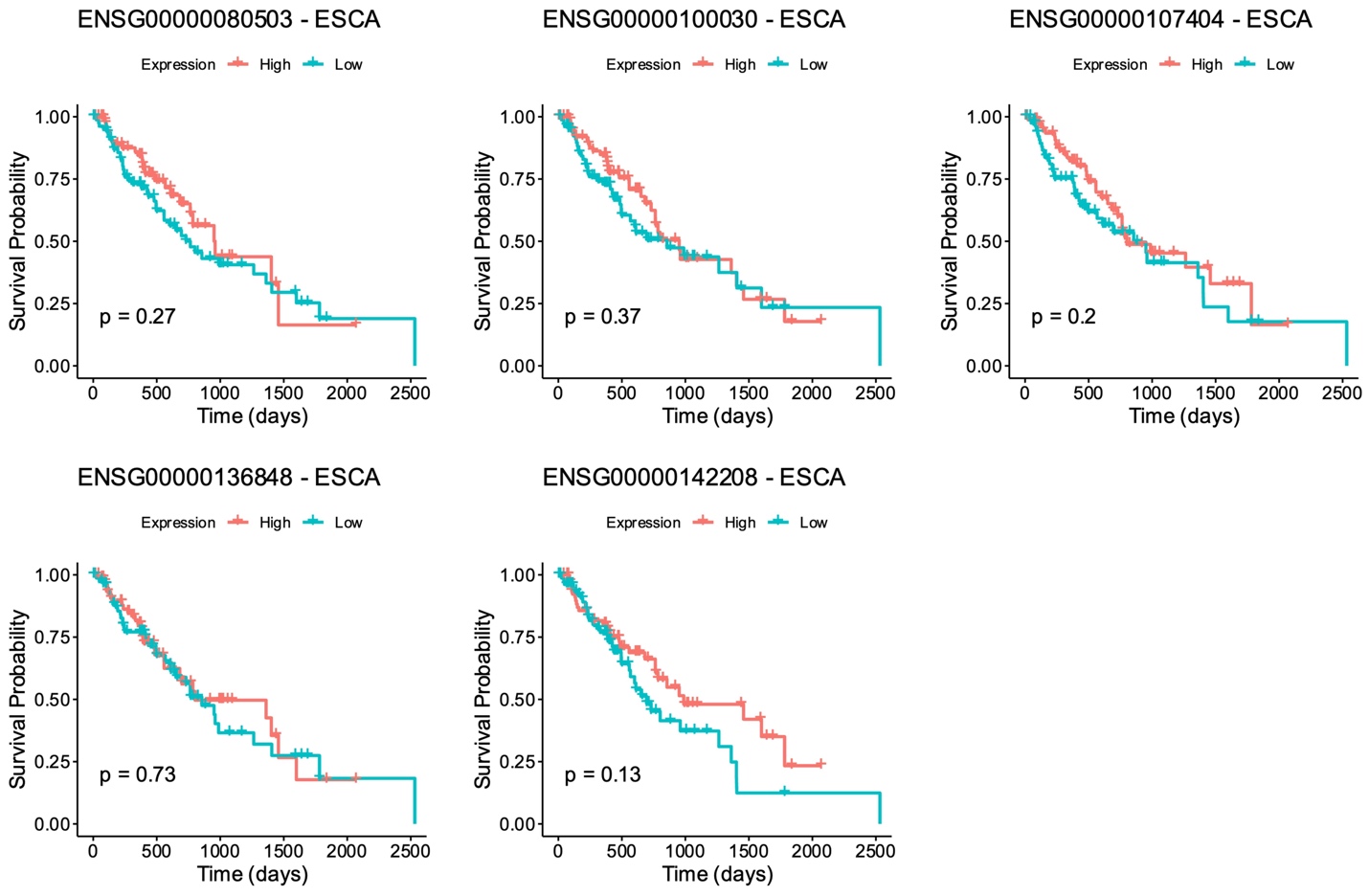

**Supplementary Figure S4.** Kaplan Meier curve of the top genes selected from cox-regression analysis in ESCA. For each gene, patients were stratified into high and low category based on median expression and p-values were estimated using log-rank test.

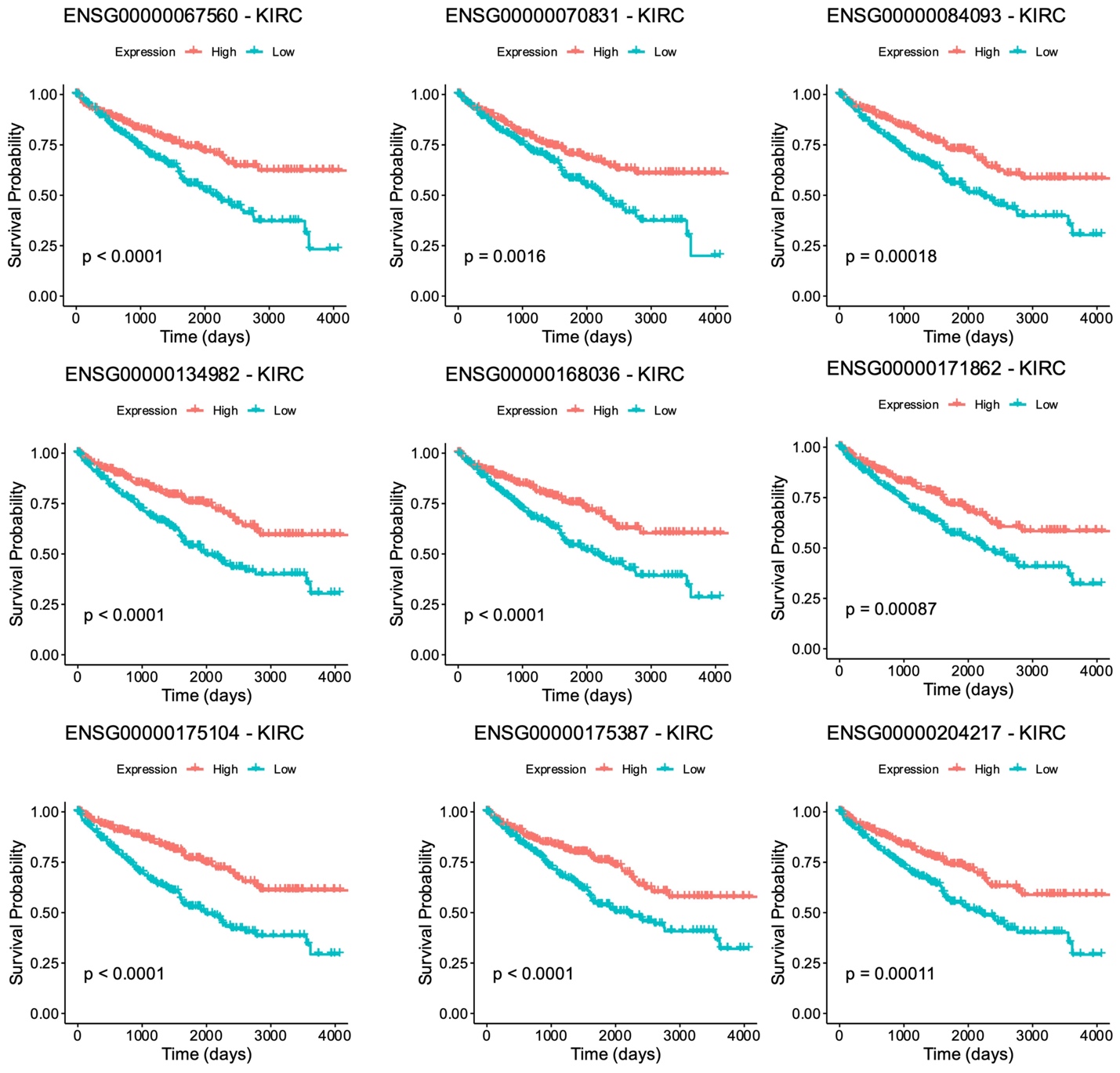

**Supplementary Figure S5.** Kaplan Meier curve of the top genes selected from cox-regression analysis in KIRC. For each gene, patients were stratified into high and low category based on median expression and p-values were estimated using log-rank test..

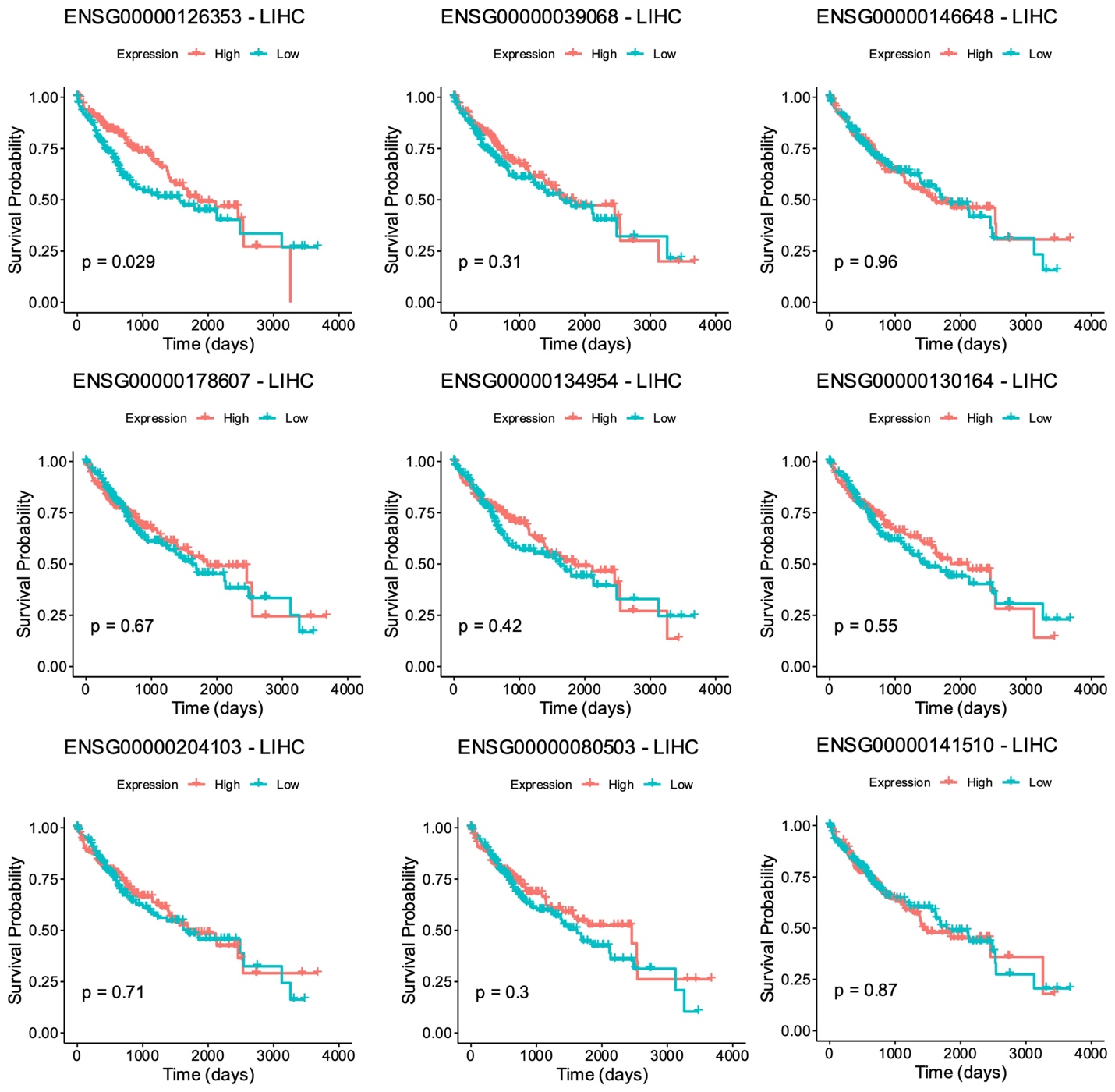

**Supplementary Figure S6.** Kaplan Meier curve of the top genes selected from cox-regression analysis in LIHC. For each gene, patients were stratified into high and low category based on median expression and p-values were estimated using log-rank test.

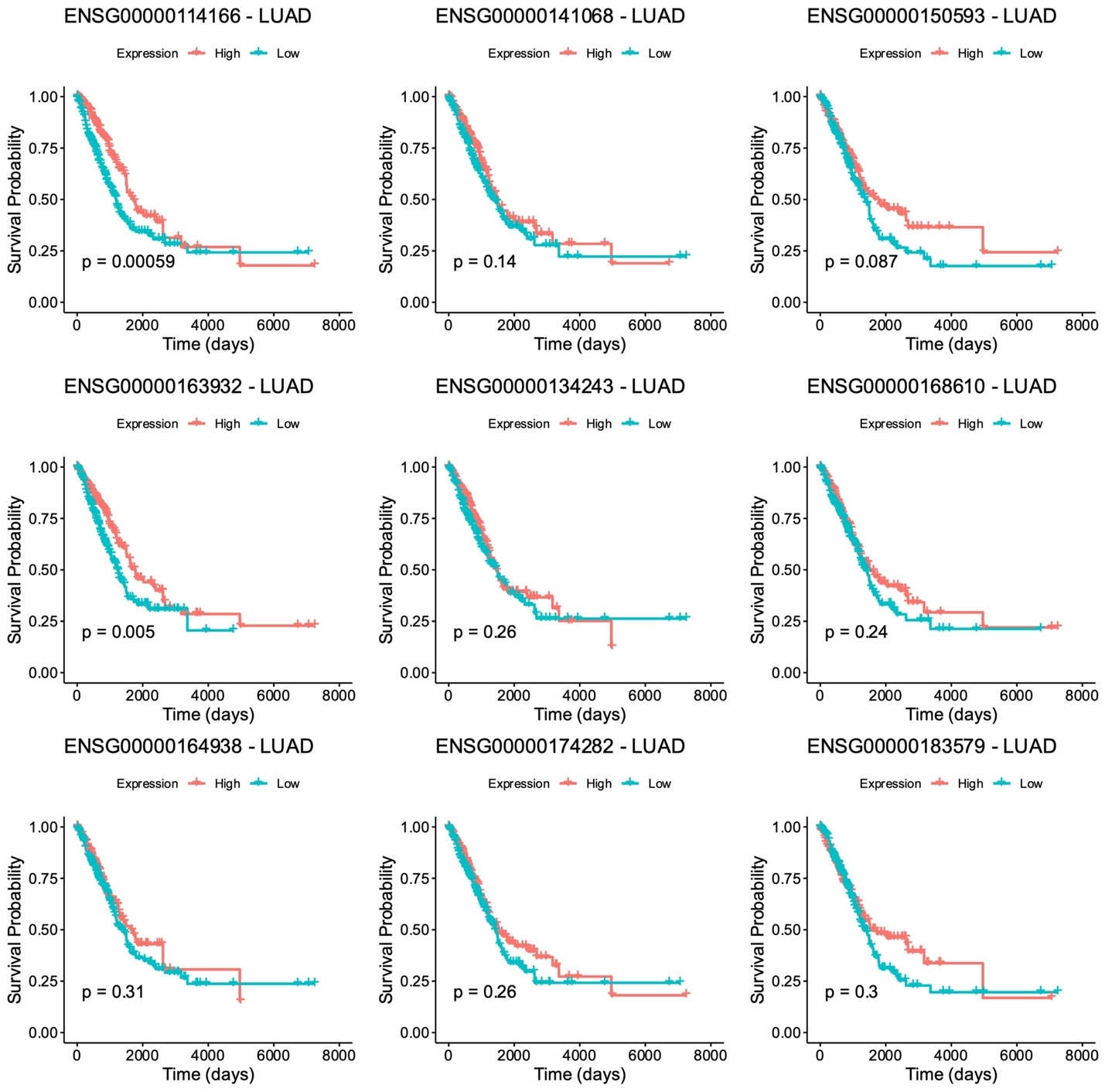

**Supplementary Figure S7.** Kaplan Meier curve of the top genes selected from cox-regression analysis in LUAD. For each gene, patients were stratified into high and low category based on median expression and p-values were estimated using log-rank test.

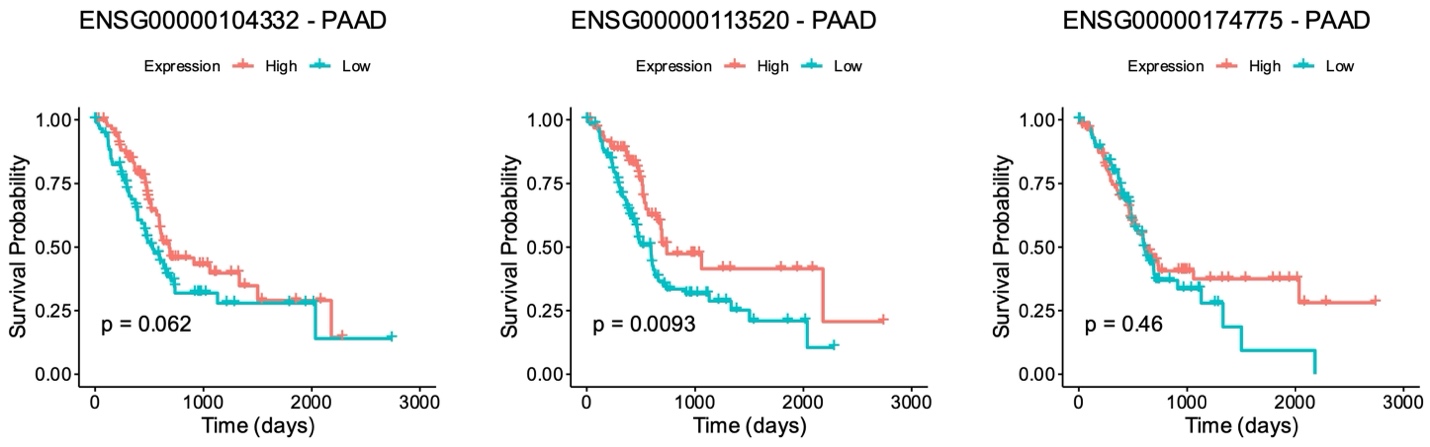

**Supplementary Figure S8.** Kaplan Meier curve of the top genes selected from cox-regression analysis in PAAD. For each gene, patients were stratified into high and low category based on median expression and p-values were estimated using log-rank test.
